# Water-Glycan Interactions Drive the SARS-CoV-2 Spike Dynamics: Insights into Glycan-Gate Control and Camouflage Mechanisms

**DOI:** 10.1101/2024.06.04.597396

**Authors:** Marharyta Blazhynska, Louis Lagardère, Chengwen Liu, Olivier Adjoua, Pengyu Ren, Jean-Philip Piquemal

**Affiliations:** LCT, Sorbonne Université, UMR 7616 CNRS, 75005 Paris, France; Department of Biomedical Engineering, The University of Texas at Austin, Texas 78712, USA Qubit Pharmaceuticals, 75014 Paris, France

## Abstract

To develop therapeutic strategies against COVID-19, we introduce a high-resolution all-atom polarizable model capturing many-body effects of protein, glycans, solvent, and membrane components in SARS-CoV-2 spike protein open and closed states. Employing *μ*s-long molecular dynamics simulations powered by high-performance cloud-computing and unsupervised density-driven adaptive sampling, we investigated the differences in bulk-solvent-glycan and protein-solvent-glycan interfaces between these states. We unraveled a sophisticated solvent-glycan polarization interaction network involving the N165/N343 residues that provide structural support for the open state and identified key water molecules that could potentially be targeted to destabilize this configuration. In the closed state, the reduced solvent polarization diminishes the overall N165/N343 dipoles, yet internal interactions and a reorganized sugar coat stabilize this state. Despite variations, our glycan-solvent accessibility analysis reveals the glycan shield capability to conserve constant interactions with the solvent, effectively camouflaging the virus from immune detection in both states. The presented insights advance our comprehension of viral pathogenesis at an atomic level, offering potential to combat COVID-19.

## Introduction

The emergence of COVID-19 pandemic (*1*) has underscored the critical importance of comprehensively understanding the biochemical nature of the severe acute respiratory syndrome coronavirus 2 (SARS-CoV-2). Despite widespread collaborative efforts within the scientific community to develop treatments and vaccines (*2, 3*), many aspects of the virus behavior remain poorly understood. Notably, elucidating the structural dynamics of the spike protein (*4–7*) is of paramount importance for discerning its functional properties and identifying viable targets for the design of new therapeutic strategies.

The coronavirus uses the spike envelope glycoproteins (S-proteins comprising subunits S1 and S2) for receptor recognition in order to bind with the host, leading to membrane fusion, as well as to the entry into the host cell to initiate the viral infection (*5, 6, 8–10*). Structurally, the spike protein features a trimeric configuration, with each protomer composed of an N-terminal domain (NTD), a receptor-binding domain (RBD), a central helix (CH), a fusion peptide (FP), a connecting domain (CD), a heptad repeat (HR), a transmembrane domain (TM), and cytoplasmic tail (CT) (Figure 1A, B) It is important to point out that the spike protein exhibits distinct prefusion conformations, namely the closed and open states (*4*). In the closed state, the spike protein is characterized by concealed RBDs (Figure 1C), therefore, limiting their interaction with the host-cell receptors. Conversely, the open state features one (Figure 1D) or multiple exposed RBDs, accessible for the binding to host cells. The transition from the closed to the open state is triggered by the binding of the spike protein to host cell receptors, such as the angiotensin-converting enzyme 2 (ACE2), allowing the viral genome to initiate its viral replication and, therefore, the infection (*9, 10*). Besides RBDs, glycans surround the SARS-CoV-2 spike protein and play a crucial role in modulating its structure and function, influencing various aspects of viral pathogenesis (*5,6,8,11–13*). While not being directly encoded in the viral genetic sequence (*14*), glycans act as a protective barrier, camouflaging the underlying protein structure from the host immune system, thus, promoting the virus infection (*12, 15, 16*). Among experimental studies on elucidating the role of spike-glycan interactions (*9, 10, 12, 13, 15–24*), the work of Huang et al. demonstrated that enzymatic removal of glycans from the SARS-CoV-2 spike protein enhances immune responses and protection in animal models, further highlighting the significance of glycans in viral pathogenesis and vaccine development (*22*).

**Fig. 1.**
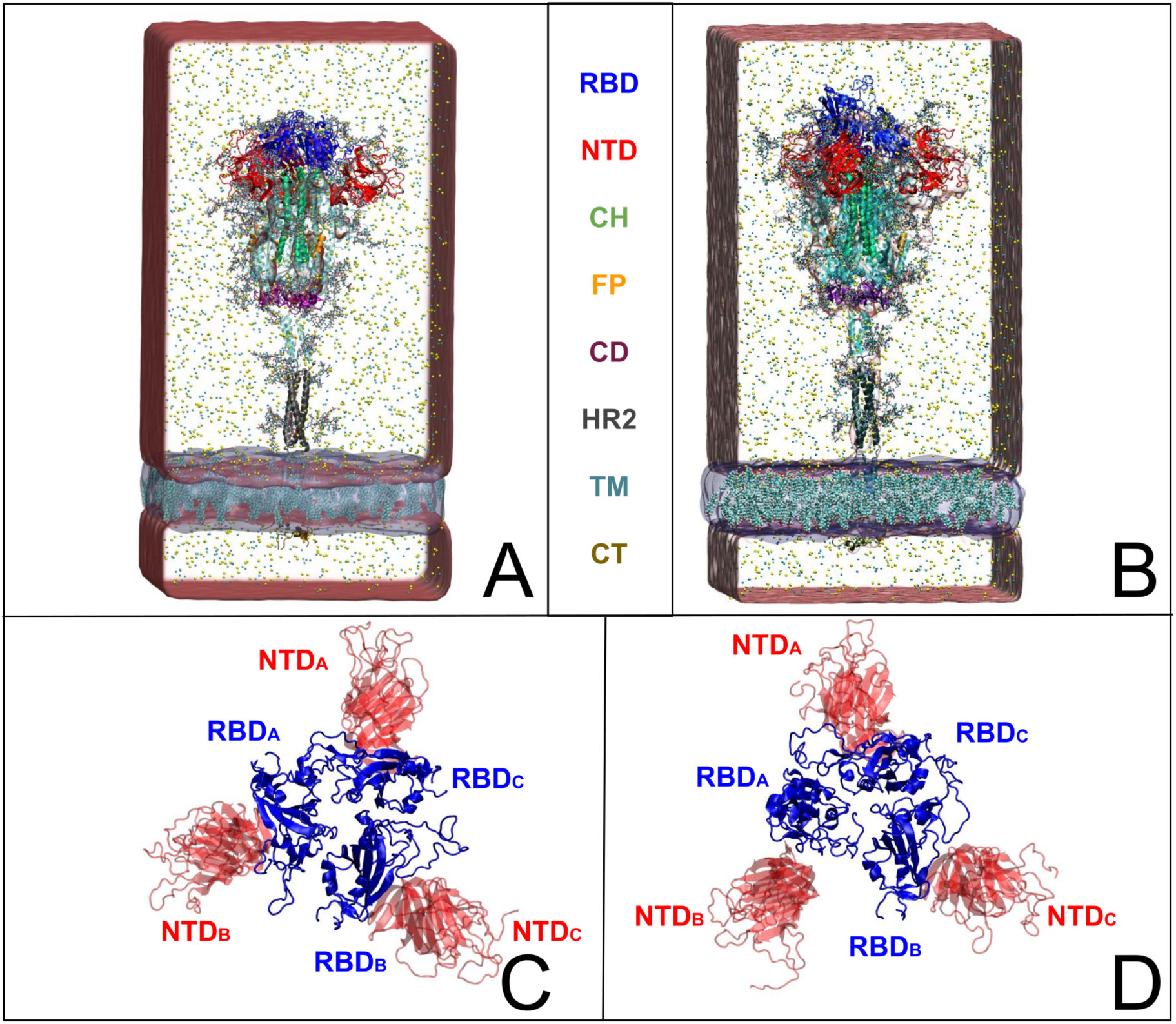
SARS-CoV-2 full spike protein structure shown in (A, C) closed and (B,D) open (one RBD_A_ up–conformation) states embedded in water box and the viral membrane with cholesterol molecules (depicted as transparent red and blue rectangles, respectively). The receptor binding domain (RBD), the N-terminal domain (NTD), the central helix (CH), the fusion peptide (FP), the connector domain (CD), the heptad repeat 2 (HR2), the transmembrane domain (TM) and the cytoplasmic tail (CT) are represented in blue, red, green, orange, mauvre, gray, cyan, and ochre, respectively. Top views (C and D) illustrate the structural reorganization of immune recognition sites on RBD and NTD between closed (C) and open (D) states.

Additionally, significant computational efforts have been dedicated to uncover intricate interactions between the spike protein and surrounding glycans (*5,6,8,25*). Thus, employing molecular dynamics (MD) simulations, Amaro et al. (*8*) showed that during the RBD opening, the glycan present at N234 undergoes an inward rotation, filling the resultant void and contributing to the stabilization of the open configuration. Subsequently, their recent MD investigation, complemented by cryo-electron microscopy and biolayer interferometry experiments (*25*) revealed that the glycans positioned at N343, N234, and N165 play a pivotal role in the spike opening by sequentially interacting with multiple residues within the RBD, a mechanism referred to as "glycan gating". It should be noted that in their studies, Amaro et al. (*8, 25*) utilized classical non-polarizable MD simulations tuning the conformational sampling to get insight on the spike opening and offering valuable insights into spike-glycan dynamics. Despite being all-atom, the non-polarizable classical MD-based approaches may not fully capture some important mechanisms of charge redistribution in complex molecular systems, therefore, potential gaps in understanding the complete spectrum of spike protein behavior should be considered. Particularly, classical MD simulations may inadequately depict weak intermolecular interactions involving many-body effects (*26*), requiring the use of polarizable force fields (*27*) for more nuanced exploration. In this context, the use of High-Performance Computing (HPC) via a distributed cloud infrastructure in conjunction with advanced sampling algorithms offers a promising avenue for the high-resolution description of intricate dynamics, especially for large and convoluted biological systems, such as SARS-CoV-2 spike protein.

To address these questions, we leveraged the Tinker-HP simulation package (*28*) employing its GPU-accelerated implementation (*29*), allowing us to conduct a series of all-atom MD simulations using the AMOEBA polarizable force field (*27, 30*). Furthermore, our simulations rely on extensive density-driven adaptive conformational sampling (*31*) of both open and closed configurations provided by Amaro et al. (*8*). It is worth noting that our study marks the first realization of a parametrization of all the components of the full-length spike protein including glycans, water solvent, counter-ions, and a membrane bilayer with a polarizable force field, thus, allowing us to access a more comprehensive and accurate representation of the spike complex interactions.

Our findings revealed subtle differences in protein-glycan interactions between the spike protein closed and open states. Notably, while the overall integrity of the glycan shield remained remarkably consistent in both forms, the mechanisms governing local interactions varied. In the open state, highly polarizable water molecules created dynamic hydrogen bonding that mediated specific protein-glycan interactions, particularly at residues N343 and N165, likely contributing to stabilizing the spike opening. In contrast, the closed state features loosely packed, low-polarization water molecules at the protein-glycan interface, suggesting that the stability of the protein structure in this state is maintained due to the inner reorganization of the protein and stabilization arising from the interacting glycans. The presented insights highlight the intricate balance between the SARS-CoV-2 state and the solvent interactions, indicating a sophisticated viral adaptation toward evading immune detection. Overall, our study provides a foundation for the development of targeted therapeutics aimed at disrupting viral infectiousness and enhancing host immune recognition, therefore, offering promising avenues for fighting COVID-19 and other viral diseases.

## Results

### Principal Component Analysis

To gain insights into the structural dynamics governing the open and closed states of SARS-CoV-2 spike protein, we employed the principal component analysis (PCA). Leveraging the solute-only trajectories shared by the Amaro laboratory (*32*) and following their PCA calculation strategy discussed in detail in the reference 8, we replicated their PCA spaces for both open and closed conformations. Subsequently, we projected our extensively sampled simulations onto such spaces (Fig. 2). It should be noted that in their work presented by Casalino et al. (*8*), the Amaro laboratory implemented an enhanced sampling strategy solely for the open state, whereas exploration of the closed state was carried out through production simulations. Despite the limited conformational sampling in the latter, our analysis revealed an emergence of multiple intersecting clusters in their simulations, indicating the conformational heterogeneity within this state (Fig. 2C). Furthermore, our extensive sampling strategy applied to the closed state (Fig. 2D) enabled the delineation of distinct clusters corresponding to both highly visited and rare conformations within this state. In contrast, a significant difference in the PCA spaces was observed in the comparison with open-state simulations. While Amaro’s data showed distinct, non-interacting clusters for this state (see Fig. 2A), our analysis depicted more intertwined clusters (Fig. 2B). It is worth mentioning that similar improvements in configuration space sampling over conventional MD have been observed previously in our group (see reference 31), thus underscoring the effectiveness of the implemented adaptive sampling strategy coupled with a polarizable force field.

**Fig. 2.**
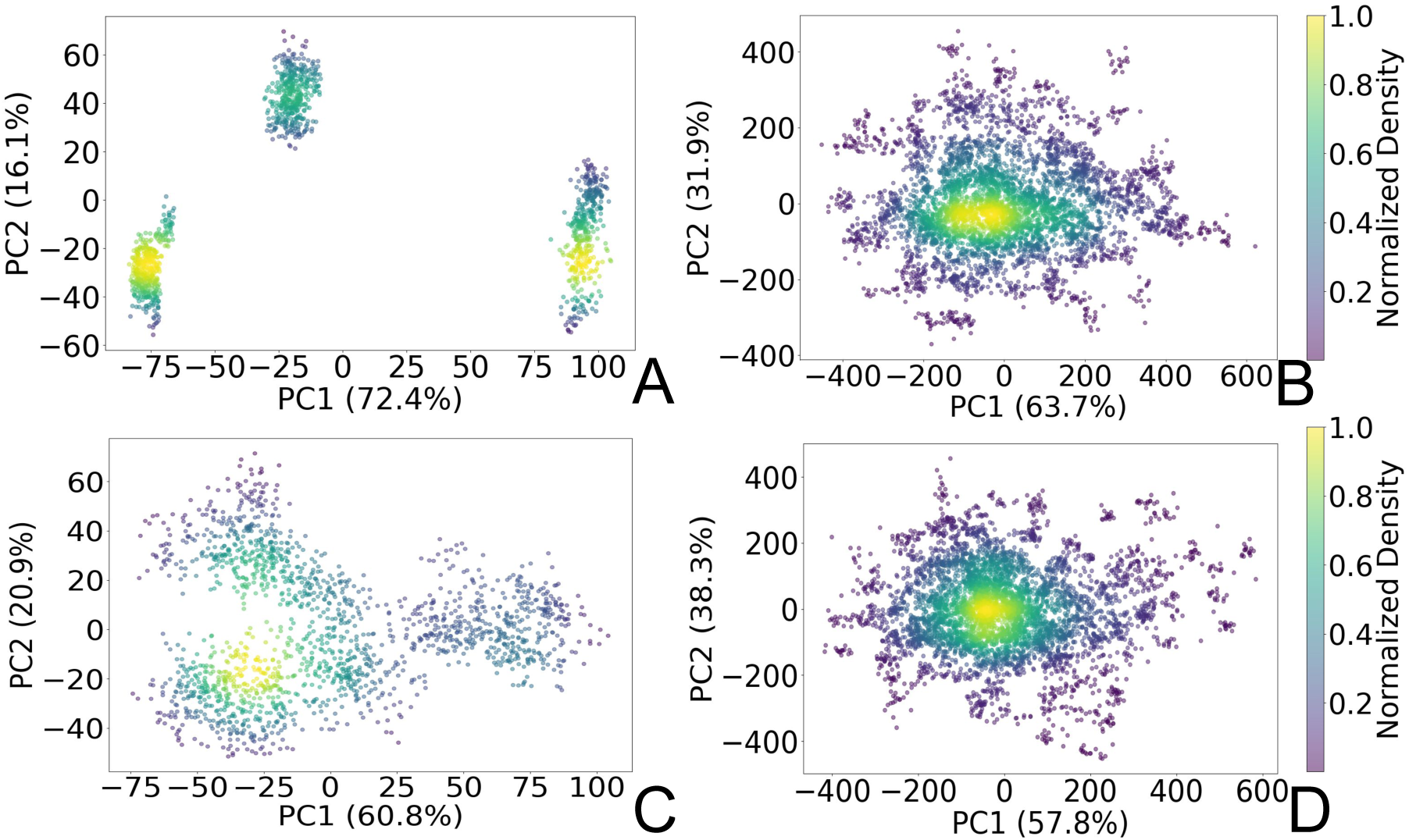
PCA plots representing two principal components (PC1 and PC2) of the SARS-CoV-2 spike C*_α_* atoms in (A) open state reproduced from the Amaro laboratory enhanced sampling simulations from three replicas, (B) open state projected to Amaro laboratory PCA space (this work), (C) closed state reproduced from Amaro laboratory production simulations from three replicas, (D) closed state projected to Amaro laboratory PCA space (this work). The points are colored based on the normalized probability densities obtained from kernel density estimation. The amount of variance of each PC is shown in %. Images are generated with Matplotlib Python library (*33*).

### Analysis of Protein and Glycans Dynamics

To validate our models against those documented by Casalino et al. (*8*), we complemented our analysis by examining the root mean square fluctuations (RMSF) of protein C*_α_* atoms elucidate the protein dynamic behavior in open and closed states (Fig. 3A). Our results revealed that the RMSF profiles derived from simulations utilizing the polarizable force field exhibit diminished fluctuations compared to the classical MD data (*8*). Such discrep-ancy likely stems from the differences in capturing the electrostatic interactions within the protein by affecting the arrangement and stability of near to C*_α_* atoms charged groups. Indeed, classical force fields, which typically employ fixed charges assigned to atoms, tend to oversimplify these interactions since they do not consider many-body polarization effects. The application of the polarizable force field provides a route to include these effects via a more accurate representation (*27, 30, 34*).

**Fig. 3.**
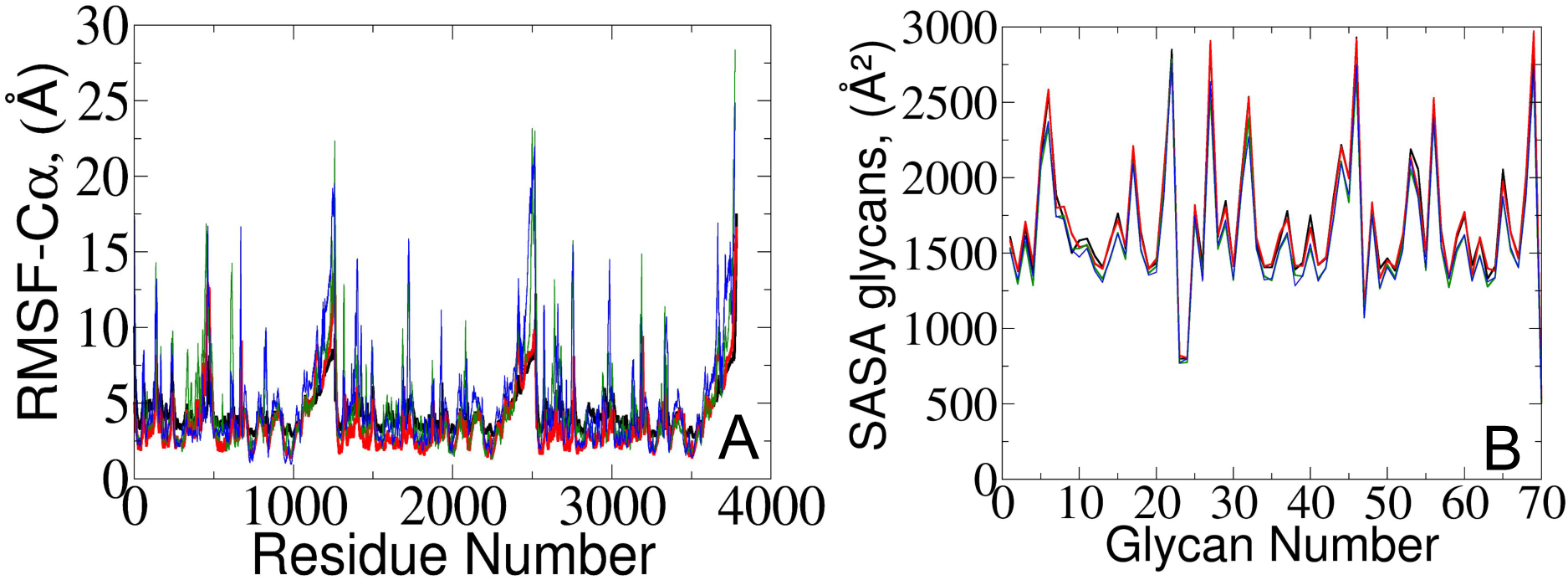
(A) RMSF of protein C*_α_* atoms and (B) averaged SASA of 70 surrounding glycans obtained from the current study (represented in red and black) and from simulations reported by Amaro lab (*8*) (green and blue) for open and closed, respectively.

Additional hydrogen-bonding and salt-bridge analysis (see Fig. S1-S6 and Data S1 in the Supplementary Materials (SM)), provides further protein structural details. From our polarizable simulations, we identified only 27 common stable inner hydrogen bonds present in both the open and closed states. The closed state exhibited 77 unique stable hydrogen bonds and 4 unique salt bridges, while the open state had 45 unique stable hydrogen bonds and only 2 unique salt bridges. Such disparity indicates that the inner protein structure of the closed state is more extensively stabilized by interactions between its residues compared to the open state, suggesting a more rigid structure, which may contribute to its structural integrity until the fusion event unfolds. Conversely, the open state, with fewer stabilizing interactions, appears more plastic and adaptable, facilitating processes such as receptor binding and viral entry.

To further gain insights into potential triggers modulating SARS-CoV-2 protein conformational changes, we investigated the average solvent-accessible surface area (SASA) of the glycans covering the protein surface. The total SASA for each glycan was accumulated across all simulation frames, and the average value was determined by dividing the total SASA by the total number of frames (Fig. 3B). Remarkably, our analysis of surrounding glycans showed similar average SASA profiles in both classical and polarizable simulations. Regardless of the conformational state and the protein structural changes along the trajectories, the glycans maintained a stable level of exposure to the surrounding solvent molecules. Such stability suggests the formation of a consistent and robust protective barrier, effectively camouflaging (*17–19,23*) the underlying protein structure from the external environment. Prior literature underscores the crucial role of glycan camouflage in CoV spike proteins (i.e, SARS-CoV-2), impacting host attachment, immune responses, and virion assembly (*11, 16–19, 35, 36*). However, it was previously unknown that the glycan covering persists unchanged across different conformational states of the spike protein, suggesting a sophisticated strategy employed by the virus to limit accessibility to the spike and potentially escape detection by the immune system.

It should be noted that despite similar glycan SASA profiles, the interaction patterns at the protein-glycan interface differ between the two states. To gain insights into glycan spatial dynamics around the spike protein, we conducted analyses of glycan root mean square deviation (RMSD) and their radial positioning relative to the protein central axis (Fig. 4A,B). The RMSD analysis revealed lower values for glycans in the closed state, indicating a more rigid and tighter packing around the protein, which likely enhances its structural integrity. Glycan radial distance analysis (Fig. 4C), along with protein-glycan dynamic cross-correlation (Fig. S7) and relative distance evaluation (Fig. S9), supported a closer association between glycans and the protein surface at immune recognition sites (i.e., the RBD and N-terminal domain (NTD)) in the closed state.

**Fig. 4.**
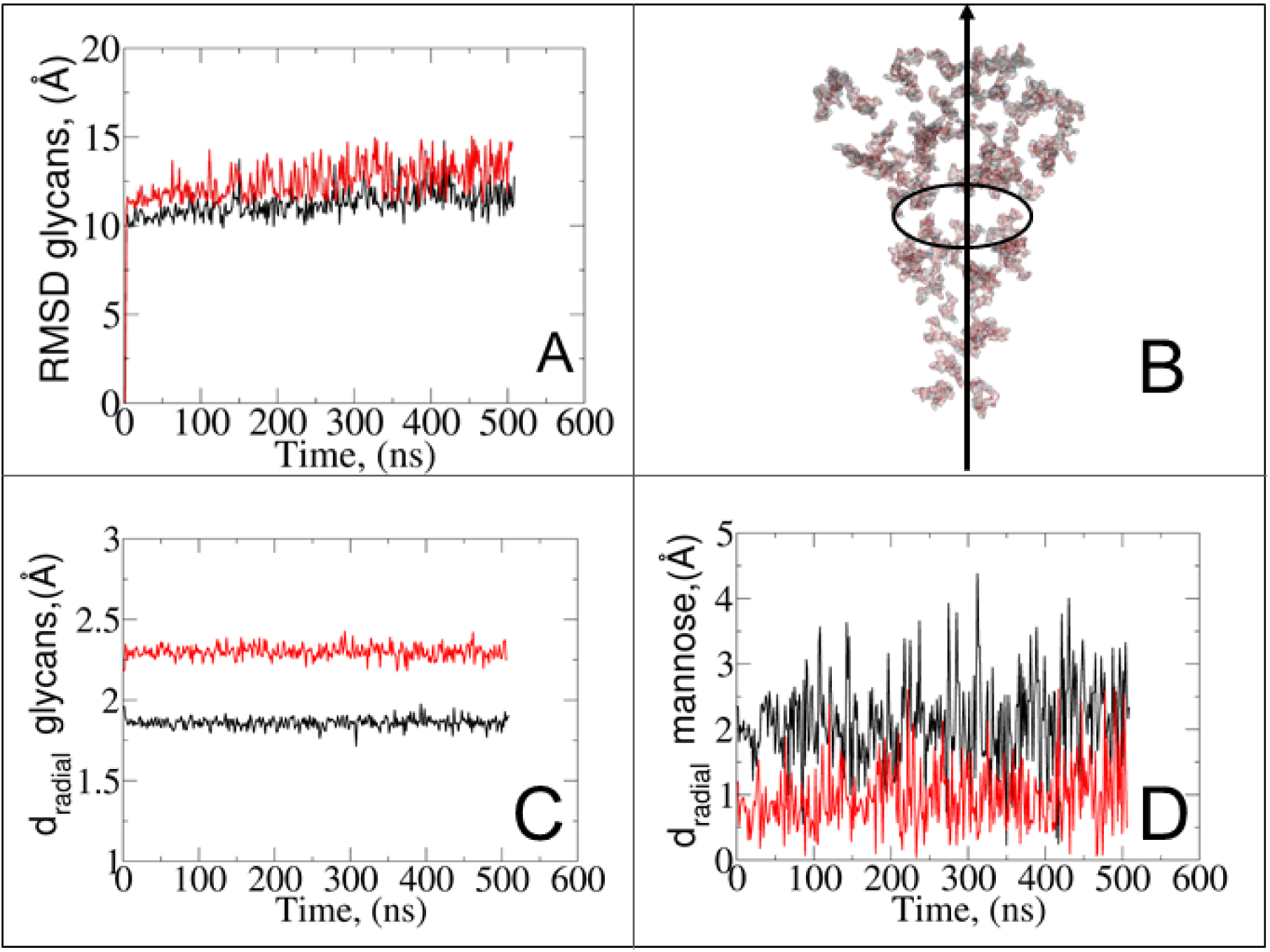
(A) RMSD of all surrounding the spike glycans excluding hydrogen atoms. (B) Schematic representation of the radial distance of glycans from the *z*-axis of the protein, which is used for the graphs in panels C and D. (C) Radial distances of all glycans relative to the protein. (D) Contribution of *α*-D-mannose residues in the total glycan radial distances. Red and black colors indicate the open and closed states, respectively.

Given the prevalence and role of *α*-D-mannose residues in viral attachment (*8, 23, 37–39*), we also evaluated their contribution to the glycan radial motion. In the closed state, the radial distances of these residues closely matched the total glycan radial distances (Fig. 4D), indicating their crucial role in the stabilization of the closed form. Conversely, in the open state, the smaller contribution of *α*-D-mannose residues (∼1 Å) to the total radial motion compared to all glycans (∼2.4 Å) suggests their lesser involvement in overall glycan dynamics. However, their closer radial arrangement in the open state may indicate their reorganization near the protein surface, potentially influencing viral infectivity, immune recognition, and evasion mechanisms.

### Solvent Interactions at the Glycan-Protein Interface through Radial Distribution Function

To explore the dynamics at the glycan-protein interface, we investigated its interactions with solvent using a radial distribution function (RDF) analysis. Initially, we computed two distinct RDFs: one between water and protein oxygen atoms, and another between water and glycan oxygen atoms. While the total intensity values of g(r) were small, our focus remained on qualitative aspects. Remarkably, the RDF profiles exhibited similarities between the open and closed states (Fig. 5A,B). We further refined the analysis by splitting the RDF between water and glycan oxygen atoms (Fig. 5B) into two parts: glycan oxygens adjacent to the protein interface (Fig. 5C) and those closer to the bulk solvent (Fig. 5D).

**Fig. 5.**
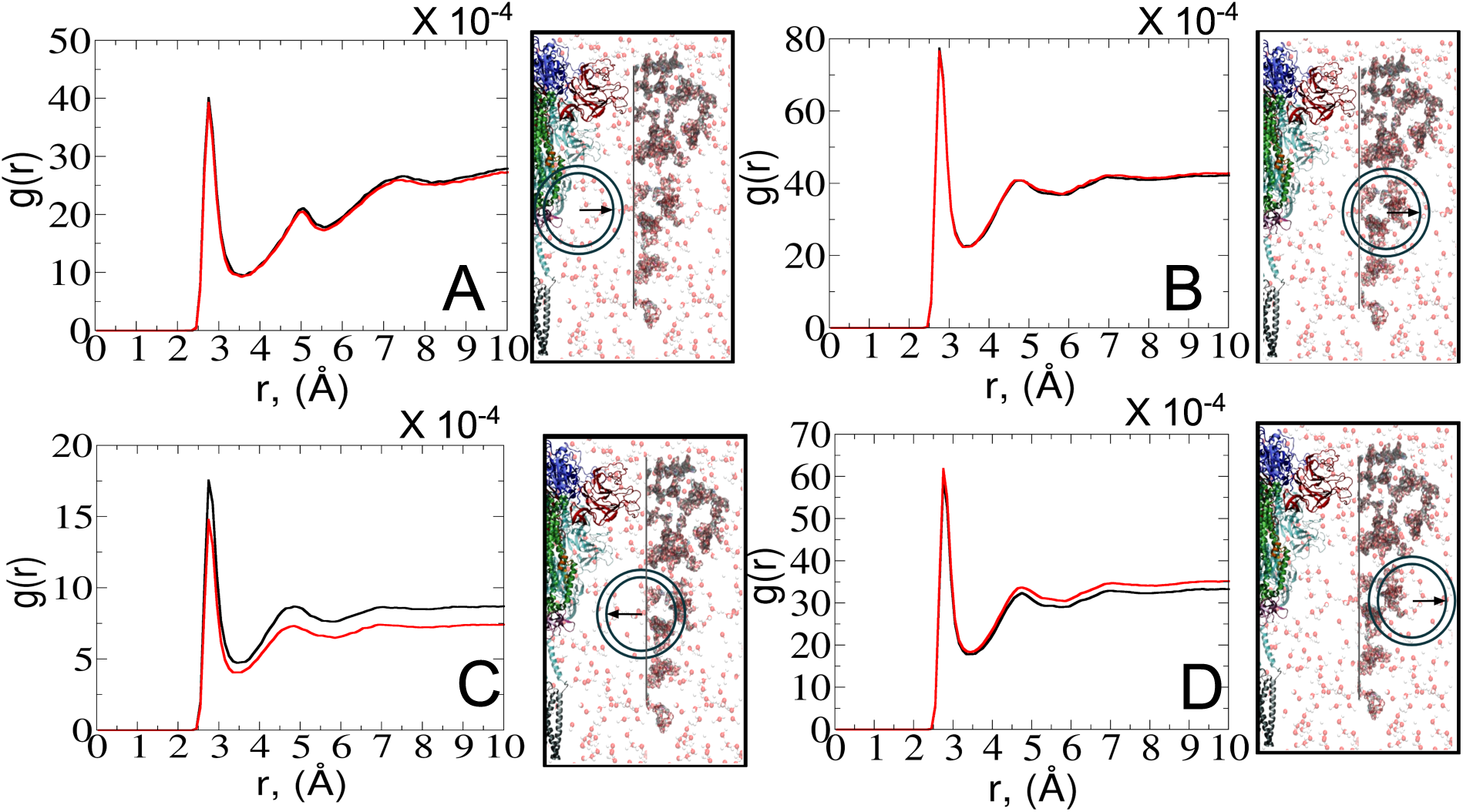
RDFs depict the interactions between water-protein and water-glycan oxygen atoms in the closed (black) and open (red) states. Each RDF plot is accompanied by a schematic representation on the right side of the graph, denoting interfaces from left to right: protein, interfacial solvent, glycan, and bulk solvent. The positions of the radial circles where the RDF calculations were performed are shown. (A) RDF between water and protein oxygen atoms. (B) RDF between water and glycan oxygen atoms. (C) RDF between water and glycan oxygen atoms adjacent to the protein interface. (D) RDF between water and glycan oxygen atoms closer to the bulk solvent.

An interesting observation emerged from the RDF of water-glycan oxygen pairs oriented towards the protein interface: the closed state exhibited a more pronounced RDF compared to the open state, indicating a potentially more permissive arrangement of interfacial water molecules. Such finding suggests that the closed conformation, characterized by a predominance of glycans near the protein surface (see Fig. S7-S9), may create a distinct microenvironment that influences the dynamics of water molecules differently compared to the open state. Conversely, the denser packing of water molecules in the open state tends to show a tighter interaction of the solvent with the protein-glycan interface, impacting its stability and functionality. The observed water behavior is further compensated by the local bulk solvent reorganization at the glycan interface (Fig. 5D).

### Role of Polarizable Water in Mediating Protein-Glycan Interactions

The disparities in water behavior at the protein-glycan interface across distinct conformational states prompted an in-depth investigation into the potential presence of solvent polarization that influences their interactions. It is important to acknowledge that only a limited number of experimental studies have explored the polarizing effect of the environment on viral activity. While investigations into enveloped viruses like SARS-CoV-2, vaccinia (*40*), and influenza (*41–43*), have shed light on the role of polarization in facilitating viral fusion processes and mediating proton transport, capturing the polarizable nature of water molecules in experiments poses significant challenges. Water dipoles, which are pivotal for polarization effects, are small in magnitude and transient, rendering their detection difficult (*44–47*). Consequently, computational investigations into solvent polarizability become essential. Non-polarizable classical force fields struggle to accurately capture the dynamic behavior of water molecules. (*27, 48*), making the use of polarizable force fields crucial for accurately representing the behavior of water molecules and elucidating their role in viral processes.

Initially, we conducted a thorough examination to identify water molecules exhibiting elevated dipole moments (i.e., surpassing 2.8 D (*49*)) and a high occupancy (i.e., exceeding 30%) proximal to the protein-glycan interface. Our analysis revealed the presence of highly polarizable water molecules at only two locations: adjacent to N343 (N343_RBD-A_) located in RBD domain in up-conformation with attached glycan G10, and adjacent to N165 (N165_NTD-B_) located in NTD (N-terminal domain) with glycan G30, exclusively in the open state (Fig. 6). Notably, these protein residues (e.g., N165 (*5, 8, 20*) and N343 (*5, 20, 21*)) have been established as pivotal players in the spike opening.

**Fig. 6.**
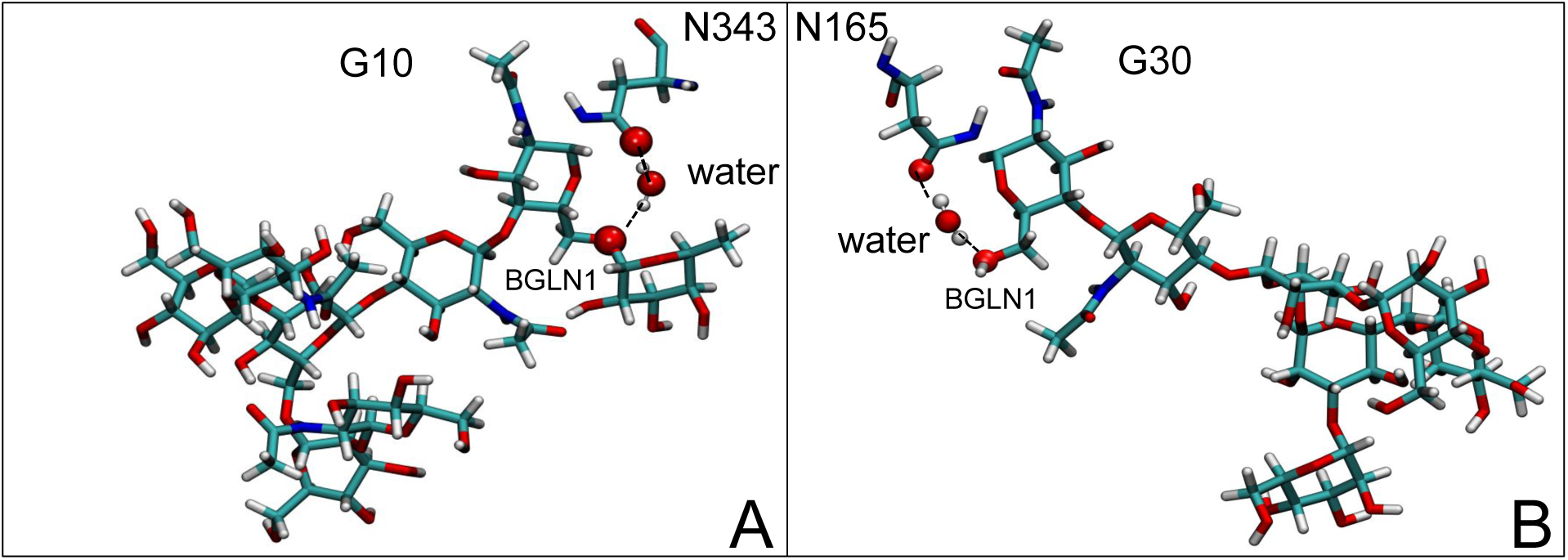
Polarizable water molecule creates bridging between (A) N343_RBD-A_ in up-conformation and *β*-N-acetyl-D-glucosamine (BGLN1) of glycan G10, and (B) N165_NTD-B_ and BGLN1 of glycan G30 of open state. In *β*-D-glucosamine, the oxygen atoms involved in this water-bridging interaction correspond to ether oxygen (A) or hydroxyl group (B). In asparagine residues, the oxygen atoms involved in this water-bridging interaction are located in the carboxamide group. Interacting patterns are shown in van der Waals representation.

Focusing on N343_RBD-A_ and N165_NTD-B_, we delved into the dynamics of their water-mediated interactions with glycans (i.e., G10 and G30, respectively). Across all iterations of the open-state simulations, we noted clear variations in water behavior at the declared locations. Benefiting from extensive conformational sampling, we hypothesized that these dynamic water fluctuations could play an integral role in the mechanism of spike opening. We identified four key phases of water behavior, based on the average dipole moments and occupancies of the polarizable water molecules across all iterations of open-state simulations. These phases encompass Bridging, Clustering, Replacement, and Relaxation, as demonstrated on the interaction interfaces of N165-G30 and N343-G10, depicted in Fig. 7.

**Fig. 7.**
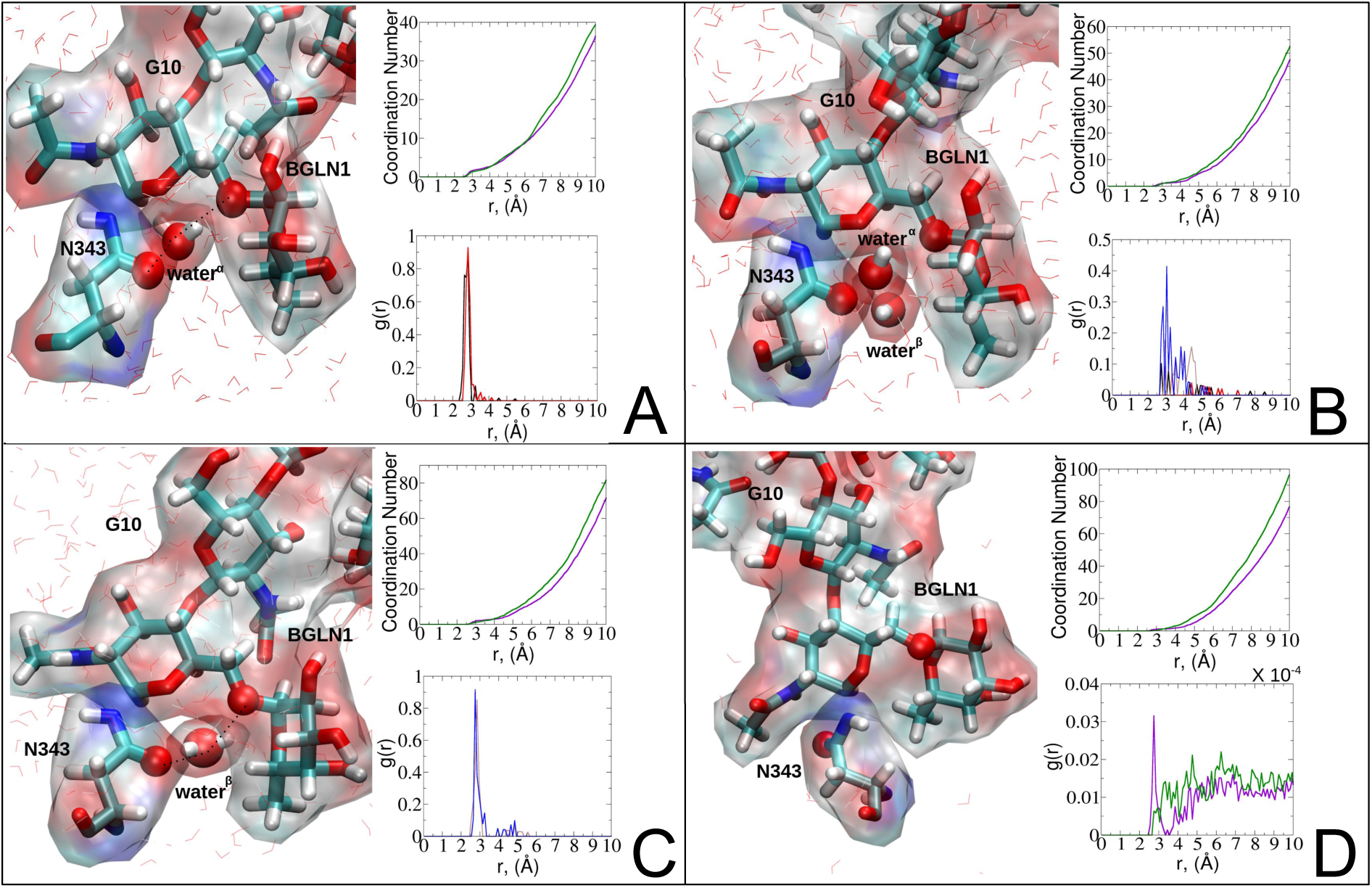
Phases of dynamic polarizable water molecule bridging formation between oxygen of N343 of RBD-A in up-conformation and oxygen of BGLN1 of glycan G10 with corresponding pair distribution functions (PDF), g(*r*), (black - N343(O)-water*^α^*(O), red - G10(O)-water*^α^*(O), brown - N343(O)-water*^β^* (O), blue - G10(O)-water*^β^* (O)), and coordination number corresponding to the interfacial water located between N343 and BGLN1 of G10 glycan residues (violet - N343(O)-water(O), green - G10(O)-water(O)). (A) Bridging through a water molecule, (B) Regrouping, (C) Replacing, (D) Relaxation. Interacting patterns are shown in van der Waals representation. To visualize the molecular surfaces of the involved moieties we employed MSMS (molecular surface) representation (*50*) available via Visual Molecular Dynamics (VMD) environment (*51*).

In the **Bridging phase**, a specific water molecule (water*^α^*) actively forms hydrogen-bond interaction between protein and glycan oxygen atoms (Fig. 7A). To be classified in this phase, water*^α^* exhibits a dipole moment that exceeds 2.8 D and an occupancy of more than 30%. To strengthen our findings, we examined the total number of water molecules proximal to the protein-glycan atoms engaged in bridging, alongside analyzing the pair distribution functions (PDFs) of water*^α^* with these atoms. The relatively high intensity of the PDF peak, nearing unity, indicates a significant concentration of water*^α^* at a distance of approximately 2.8 Å from the N343 and G10 oxygen atoms, corresponding to the first coordination number, underscoring a strong correlation between the number of water molecules present at that distance and the peak intensity of the PDF.

Conversely, during the **Clustering phase**, water*^α^* experiences a reduction in dipole moment, falling below the threshold of 2.8 D, and a decrease in occupancy, dipping below 30%, as it conglomerates with other polarizable water molecules, thereby diminishing its individual contribution to interactions. Furthermore, within this phase, another water molecule, water*^β^* , exhibits a heightened dipole moment (more than 2.8 D), positioning to potentially replace water*^α^* (Fig. 7B). Herein, the PDFs of water*^α^* and water*^β^* oxygen atoms appear notably noisy and less intensive compared to the Bridging phase, indicating a competition between these water atoms.

During the **Replacement phase**, water*^α^* is finally substituted by a highly polarizable water molecule, water*^β^* that exhibits a dipole moment exceeding 2.8 D and an occupancy of more than 30%, ensuring the continuity of interactions (Fig. 7C). During this phase, water*^β^* undergoes an initial stabilization period, reflected in the PDF displaying peaks of low intensity of 4-5 Å. Subsequently, upon achieving stability, water*^β^* facilitates a bridging interaction between the protein and glycan residues, leading to a PDF closely resembling that observed during the Bridging phase, characterized by high intensity.

Finally, the **Relaxation phase** signifies the absence of significant water interactions at the interface (Fig. 7D). Notably, comprehensive RDF functions for all water molecules proximal to the protein-glycan interface affirm the lack of pronounced interactions during this phase, with the closest interfacial water oxygen observed to be ∼ 3.5 Å. The loss of water mediation is attributed to the reorganization of the involved protein interacting sites, leading to changes in conformation and potentially disrupting the sites of interaction with water molecules.

To evaluate variations in the local environment and interactions experienced by the water molecules, we performed an in-depth analysis of the average dipole moments observed during each phase, along with their respective duration, across all iterations (Table 1). To compare, we also estimated the average dipole moments relative to the same group of oxygen atoms in the closed state, where no water-mediated interactions were captured. Additionally, we conducted a distribution analysis of the dipole moments for both the open and closed states separately (see Fig. S11-S14 in SM). The water molecules involved in bridging interactions demonstrated higher polarity compared to other interfacial water molecules located in the proximity, thereby, enhancing the stability of these interactions. As it can be noticed from Table 1, substantial variances are observed in the dipole moments of protein residues across different conformational states, indicating significant conformational changes occurring throughout the simulations, reflective of dynamic structural transitions within the protein.

**Table 1.**
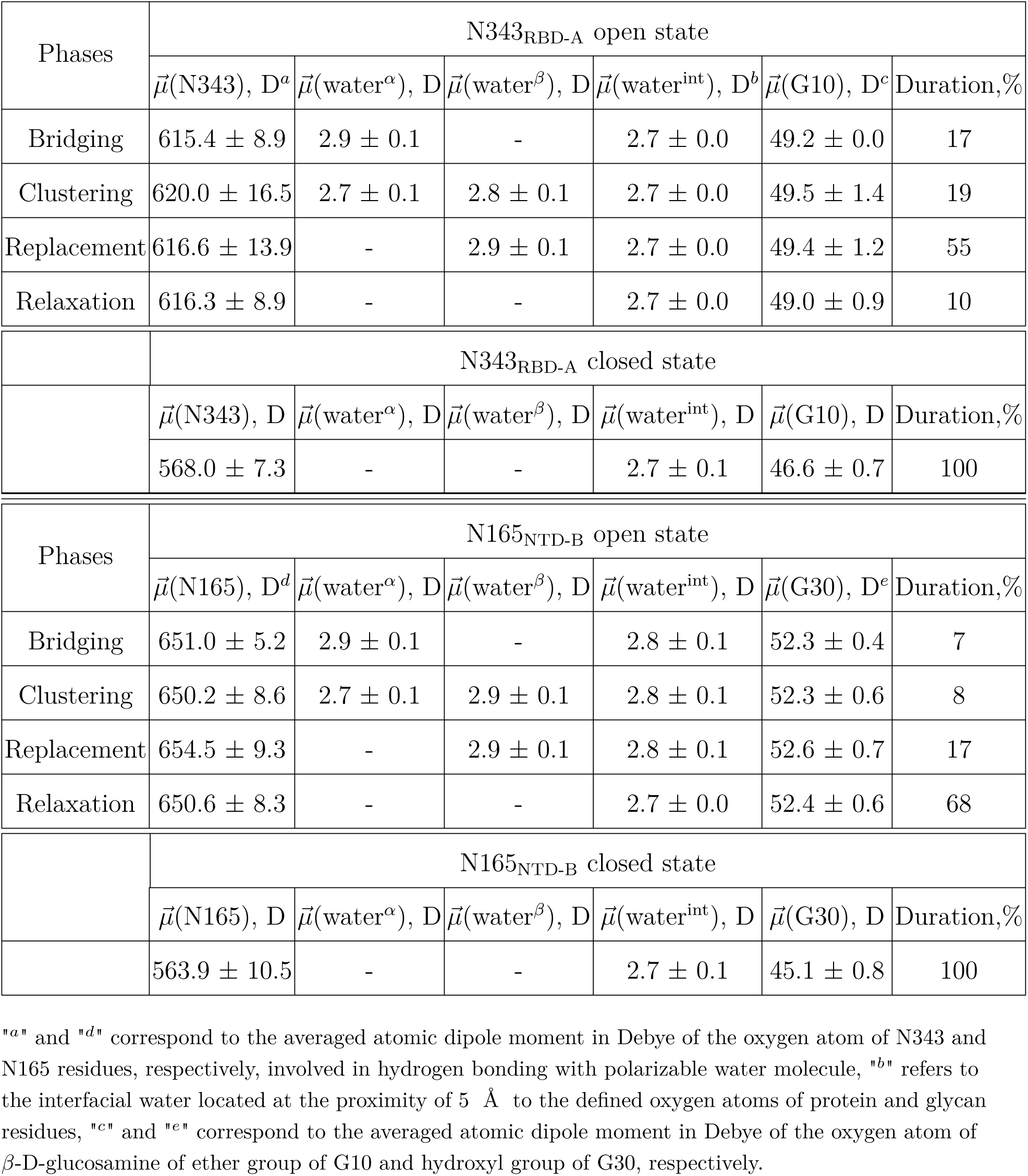
Comparison of average dipole moments and duration of water-mediated interactions in open and closed states for N343_RBD-A_ and N165_NTD-B_ residues.

| Phases | N343<sub>RBD-A</sub> open state |  |  |  |  |  |
| --- | --- | --- | --- | --- | --- | --- |
| | $\vec{\mu}(\text{N343}), \text{D}^a$ | $\vec{\mu}(\text{water}^\alpha), \text{D}$ | $\vec{\mu}(\text{water}^\beta), \text{D}$ | $\vec{\mu}(\text{water}^{\text{int}}), \text{D}^b$ | $\vec{\mu}(\text{G10}), \text{D}^c$ | Duration, % |
| Bridging | $615.4 \pm 8.9$ | $2.9 \pm 0.1$ | - | $2.7 \pm 0.0$ | $49.2 \pm 0.0$ | 17 |
| Clustering | $620.0 \pm 16.5$ | $2.7 \pm 0.1$ | $2.8 \pm 0.1$ | $2.7 \pm 0.0$ | $49.5 \pm 1.4$ | 19 |
| Replacement | $616.6 \pm 13.9$ | - | $2.9 \pm 0.1$ | $2.7 \pm 0.0$ | $49.4 \pm 1.2$ | 55 |
| Relaxation | $616.3 \pm 8.9$ | - | - | $2.7 \pm 0.0$ | $49.0 \pm 0.9$ | 10 |
|  | N343<sub>RBD-A</sub> closed state |  |  |  |  |  |
| | $\vec{\mu}(\text{N343}), \text{D}$ | $\vec{\mu}(\text{water}^\alpha), \text{D}$ | $\vec{\mu}(\text{water}^\beta), \text{D}$ | $\vec{\mu}(\text{water}^{\text{int}}), \text{D}$ | $\vec{\mu}(\text{G10}), \text{D}$ | Duration, % |
| | $568.0 \pm 7.3$ | - | - | $2.7 \pm 0.1$ | $46.6 \pm 0.7$ | 100 |
| Phases | N165<sub>NTD-B</sub> open state |  |  |  |  |  |
| | $\vec{\mu}(\text{N165}), \text{D}^d$ | $\vec{\mu}(\text{water}^\alpha), \text{D}$ | $\vec{\mu}(\text{water}^\beta), \text{D}$ | $\vec{\mu}(\text{water}^{\text{int}}), \text{D}$ | $\vec{\mu}(\text{G30}), \text{D}^e$ | Duration, % |
| Bridging | $651.0 \pm 5.2$ | $2.9 \pm 0.1$ | - | $2.8 \pm 0.1$ | $52.3 \pm 0.4$ | 7 |
| Clustering | $650.2 \pm 8.6$ | $2.7 \pm 0.1$ | $2.9 \pm 0.1$ | $2.8 \pm 0.1$ | $52.3 \pm 0.6$ | 8 |
| Replacement | $654.5 \pm 9.3$ | - | $2.9 \pm 0.1$ | $2.8 \pm 0.1$ | $52.6 \pm 0.7$ | 17 |
| Relaxation | $650.6 \pm 8.3$ | - | - | $2.7 \pm 0.0$ | $52.4 \pm 0.6$ | 68 |
|  | N165<sub>NTD-B</sub> closed state |  |  |  |  |  |
| | $\vec{\mu}(\text{N165}), \text{D}$ | $\vec{\mu}(\text{water}^\alpha), \text{D}$ | $\vec{\mu}(\text{water}^\beta), \text{D}$ | $\vec{\mu}(\text{water}^{\text{int}}), \text{D}$ | $\vec{\mu}(\text{G30}), \text{D}$ | Duration, % |
| | $563.9 \pm 10.5$ | - | - | $2.7 \pm 0.1$ | $45.1 \pm 0.8$ | 100 |

"a" and "d" correspond to the averaged atomic dipole moment in Debye of the oxygen atom of N343 and N165 residues, respectively, involved in hydrogen bonding with polarizable water molecule, "b" refers to the interfacial water located at the proximity of 5 Å to the defined oxygen atoms of protein and glycan residues, "c" and "e" correspond to the averaged atomic dipole moment in Debye of the oxygen atom of β-D-glucosamine of ether group of G10 and hydroxyl group of G30, respectively.

Besides, a net contrast arises from the comparison between the absolute values of the dipole moments of the oxygen of the carboxamide group of glycan-gating residues N343 and N165 in the open state, which could be attributed to the differences in their local environments at the protein-solvent-glycan interface and functional roles within the protein structure. Specifically, N343 experiences a more pronounced interaction with surrounding water molecules, which is also reflected in the cumulative duration of the Bridging and Replacement phases (exceeding 70%), leading to higher dipole moments compared to N165. In contrast, the absolute values of dipole moments for both N343 and N165 oxygen atoms in the closed state remain similar, suggesting that these residues may experience comparable degrees of involvement in stabilizing the protein structure in this particular state. Moreover, in the case of G30, the oxygen atom involved in water bridging is located in the hydroxyl group. The absence of high solvent polarizing effects in the closed state may result in fewer interactions contributing to the overall dipole moment, consequently, yielding a lower value compared to the open state where such interactions are more prevalent. It should be noted that the dipole moments related to the oxygen atoms of glycans G10 and G30 exhibit relatively low variances, underscoring the consistent role of glycans in shielding protein interfaces in both states, contributing to the overall stability and function of the spike.

## Discussion

In this study, we performed extensive MD simulations of two distinct prefusion SARS-oV-2 spike protein states (i.e., open and closed) using a polarizable force field coupled with advanced conformational sampling techniques. Such a combination is essential for unraveling the complex structural dynamics and molecular mechanisms of the SARS-CoV-2 spike protein, which are paramount for understanding its role in viral infectivity and identifying potential therapeutic targets against COVID-19. Through our computational approach utilizing the AMOEBA polarizable force field and extensive density-driven conformational sampling, we rigorously explored the conformational dynamics of the spike protein in both open and closed states, aggregating 1.2 *μ*s of simulation. By projecting our data onto the principal component analysis (PCA) space generated by Casalino et al. (*8*), we demonstrated the sampling capacity of our strategy (Fig. 2A,C). Our analysis unveiled stable structural states of the protein (Fig. 2B,D), characterized by central regions of high probability densities (0.6-1.0), alongside capturing rare conformations of lower densities (0-0.4), providing valuable insights into the spike dynamic behavior. Additionally, further validations, by means of root mean square fluctuation (RMSF) of the protein C*_α_* atoms (Fig. 3A), reaffirmed the reliability and accuracy of employing a polarizable force field and emphasized the importance of an accurate representation of the electrostatic interactions for understanding the viral protein behavior of SARS-CoV-2. Given that the spike is a glycoprotein, a key question arises regarding the role of the surrounding glycan oligosaccharides in modulating its structural dynamics and functional properties. To address this question, we thoroughly examined the glycan solvent-accessible surface area (SASA) in both states (Fig. 3B). Remarkably, our analysis revealed a consistent pattern: regardless of the spike conformational state, the glycans exhibited uniform levels of exposure to the solvent, forming a stable glycan shield. The striking stability in glycan exposure to the solvent suggests a sophisticated strategy employed by the virus to control access to the protein surface. In terms of viral activity, such uniform glycan behavior tends to indicate the mechanism of adeptness of the virus at evading immune detection, enabling its persistence and spread within the host regardless of its conformational state. Additional analysis (Fig. S1-S9 in SM) of the interaction patterns within the protein network revealed noticeable differences between the open and closed states, particularly in hydrogen bonding and salt bridge formations. Specifically, the closed state exhibited more stable interactions, indicating a more rigid structure that likely contributes to maintaining integrity until the fusion event. Further examination of the glycan shield flexibility and radial motion with respect to the protein surface unraveled a more compact and stiff glycan association to the critical protein immune recognition sites (i.e., RBD and NTD) in the closed state, suggesting their role in maintaining the structural integrity of the closed form. Additionally, the crucial role of *α*-D-mannose residues in stabilizing the closed conformation is evidenced by their radial distance alignment with overall radial glycan distances. In contrast, their closer reorganization to the protein surface of the open state suggests a nuanced role in modulating viral infectivity. Our observation aligns with recent studies (*52–62*) that explored lectin-based inhibitors targeting high-mannose residues on coronaviruses, suggesting a viable strategy for preventing viral entry and enhancing immune recognition.

Recognizing that the virus may fine-tune its infectivity and immune evasion strategies, we further sought to gain insights into how the solvent dynamics influences protein-glycan interactions and, consequently, the stability of the glycan shield. To achieve this, we analyzed the dynamical interactions occurring in this region by means of radial distribution function (RDF) (Fig. 5). While overall RDF profiles for water molecules in proximity to the protein-glycan surface appeared similar between the open and closed states, mirroring the consistent SASA patterns (Fig. 3B), a detailed examination revealed differences at the water-glycan interface. In particular, the water molecules surrounding the glycans displayed a more dispersed arrangement around the protein-glycan interface in the closed state, potentially influencing the protein conformational stability and functional properties differently from the open state. The observed arrangement of interfacial water molecules prompts consideration that the mechanism underlying the maintenance of closed or open conformation may differ, underscoring the intricate adaptability of the virus to varying environmental conditions.

It should be noted that the use of the spike polarizable model provides a unique opportunity to delve into the many-body effects induced by the solvent. Based on the differences in the interfacial water behavior along the open and closed states, we decided to investigate whether the polarizable nature of water molecules influences the stability and dynamics of protein-glycan interactions. Our investigation revealed that highly polarizable water molecules, exclusively present in the open state, play a crucial role in mediating interactions between specific residues, notably N343 and N165 (*5, 20, 21, 25, 63*) (Fig. 6), and their corresponding glycans (i.e., G10, G30), shedding light on potential mechanisms underlying spike protein opening. In this context, it is important to highlight the critical discovery of glycan gating by Sztain et al. (*25*). While their work emphasized the importance of glycans at N343 in initiating and N234 and N165 in supporting the RBD opening, (*25,63*) our study underscores the multifaceted nature of spike protein dynamics, integrating both glycan-mediated mechanisms and solvent interactions. Specifically, for N234, we observed only the reorganization of the attached highly-mannose glycan G31, as depicted in contact maps comparing the open and closed states (see Fig. S9). Such behavior stands in contrast to the solvent-mediated interactions involving N343 and N165 patterns.

To elucidate the water-mediating mechanism, we identified four distinct phases along the accumulated open-state trajectories: Bridging, Clustering, Replacement, and Relaxation (see Fig. 7, Table 1, and Fig. S11-S14). During the Bridging phase, a specific water molecule (water*^α^*) actively forms hydrogen bonding between protein and glycan atoms, exhibiting elevated dipole moments and high occupancy. Subsequently, in the Clustering phase, the water*^α^* concurs with another water molecule water*^β^* to form the bridge, re-sulting in the reduction in its dipole moment and occupancy. Notably, replacement by a highly polarizable water*^β^* occurs during the Replacement phase, ensuring the continuity of interactions. Finally, the Relaxation phase signifies the absence of significant water interactions at the interface, potentially due to protein conformational changes disrupting the sites of interaction. Remarkably, in the closed state, we observed only water molecules with low dipole moments in the proximity of the protein-glycan interface. Such finding aligns with the earlier observation of sparse water density around glycans in the RDF analysis (Fig. 5C), suggesting that the stability of the closed state is likely maintained due to the internal reorganization of the protein and glycan interfaces rather than the influence of the solvent.

Overall, our study underscores the viral adaptability of the SARS-CoV-2 spike protein to different conditions. Despite dynamical changes in solvent and local protein-glycan interactions, the overarching protective function of the glycan shield remains robust, ensuring viral persistence within the host. By targeting the mechanisms underlying glycan camouflaging, novel therapeutic strategies could be developed. For instance, disrupting the glycan shield could compromise the spike protective camouflaging, therefore, enhancing immune detection. Besides, the observed differences in solvent dynamics between open and closed states provide valuable insights into the spike protein opening mechanism. Targeting on the highly polarizable water-mediated interactions at the N343/N165 sites could destabilize the open state, inhibiting its interaction with ACE2. These approaches offer promising avenues for preventing viral entry and improving host immune recognition, thus providing effective strategies for combating coronavirus infection.

While our study sheds light on the structural dynamics and molecular mechanisms of the SARS-CoV-2 spike protein, it is essential to acknowledge certain limitations. Firstly, our computational approach focused solely on modeling the spike protein, which, with a total of 1,693,134 atoms, presented significant computational challenges. Despite leveraging features such as HPC, modeling the entire virus would introduce even greater complexity, computational demands, and costs. Furthermore, our analysis primarily examines structural dynamics and interactions, neglecting other important factors such as post-translational modifications and environmental influences, which could also play significant roles in viral infectivity. Additionally, our models did not incorporate the ACE2 domain, which is the receptor for the spike protein, and could provide further insights into the binding dynamics and potential changes in solvent polarization at the glycan-protein interfaces. Besides, experimental validation of our computational predictions within advanced experimental techniques capable of capturing the behavior of polarizable solvent at the glycan-protein interfaces remains crucial to confirm the observed structural dynamics and interactions.

## Materials and Methods

### Computational Assays

The starting structure files of open and closed states (6VSB (*8, 64*) and 6VXX (*8, 65*) Protein Data Bank (PDB) ids, respectively) were taken from Amaro Lab site (*32*), comprising the full-length spike protein (58,634 atoms) enveloped with 70 glycans (14,236 atoms), membrane of POPC (1-palmitoyl-2-oleoyl-sn-glycero-3-phosphocholine): POPI (1-palmitoyl-2-oleoyl-sn-glycero-3-phosphoinositol): POPS (1-palmitoyl-2-oleoyl-sn-glycero-3-phosphatidylserine): POPE (1-palmitoyl-2-oleoyl-phosphatidylethanolamine) and cholesterol (in total of 165,743 atoms), water molecules (1,451,508 atoms), sodium (1,647) and chloride ions (1366) creating a physiological concentration of 150mM. The total rectangular box size was 204.7 × 199.4 × 408.5 Å^3^. The protein, water, and counterions were parameterized by AMOEBA polarizable force field by means of **PDBXYZ** utility avail-able in the TINKER software package. More guidelines about the TINKER source code can be found in reference 66. The AMOEBA parameters for the rest of the system (i.e. lipids and glycans) were generated within Poltype2, created for automated parametrization of small molecules (for more parametrization details see Data S2 in SM). Additional details about the Poltype2 parametrization tool are available in the public GitHub repository (*67*) and in references 68, 69. The final structure files along with the parametrization files are provided in Data S3-S4 in SM.

### Advanced Molecular Dynamics Simulation Details

To conduct an extensive unsupervised adaptive sampling simulation of open and closed states, we initiated our study with 10-ns conventional MD simulations employing the AMOEBA polarizable force field. Each system underwent an initial minimization process using a limited-memory Broyden–Fletcher–Goldfarb–Shanno (L-BFGS) algorithm until achieving a Root Mean Square (RMS) gradient of 1 kcal/mol. Satisfied with the resultant models for both states, we proceeded with the extensive adaptive sampling.

For each MD simulation, we utilized the BAOAB-RESPA1 integrator (*70*) implemented in the Tinker-HP software package (*70*), which is available for public usage in GitHub repository (*71*) or in the Tinker-HP Web site (*72*). The integration scheme incorporated two time-step sizes: a short time step of 0.001 ps and an intermediate time step of 0.004 ps. The short time step was used to accurately capture the fastest motions within the system, while the intermediate time step addressed medium-frequency motions and interactions.

Our sampling technique, elaborately described in the reference 31, is a multi-iterative approach tailored for execution on large supercomputers equipped with hundreds of GPU cards. Initially, we conducted nine independent 10-ns simulations, each starting from the same conformations for both states but with varied initial velocities. Subsequently, the resulting structures from these simulations were aligned with those obtained from the study by Casalino et al. (*8*). Further fully automated structure selections were made by means of principal component analysis (PCA) based on their density in a low-dimensional space, prioritizing exploration of less-explored regions. Multiple 0.5-ns simulations were then launched from these initial structures, with each simulation state recorded at regular intervals. Cumulatively, 340 micro-iterations were performed to obtain a total simulation time of 510 ns for each state, allowing to capture a significantly broader conformational landscape compared to traditional single trajectory sampling methods. It should be noted that in each iteration, we computed a debiasing score to evaluate observation significance, and therefore, ensuring a more accurate depiction of the conformational landscape. Table 2 provides detailed information about the adaptive sampling process, including the number of micro-iterations, the duration of each micro-iteration, and the resulting speed of the simulations.

**Table 2.** Details of the adaptive sampling process.

| # of macro-iteration | # of micro-iterations | Length, ns/micro-iteration | Speed, ns/day |
| --- | --- | --- | --- |
| 1 | 10 | 0.51 | 1.5 |
| 2 | 25 |  |  |
| 3 | 25 |  |  |
| 4 | 50 |  |  |
| 5 | 50 |  |  |
| 6 | 45 |  |  |
| 7 | 45 |  |  |
| 8 | 45 |  |  |
| 9 | 45 |  |  |

### Adaptive Sampling on the Cloud

In recent years, cloud computing has emerged as an invaluable resource for handling computing demands, with providers such as Amazon Web Services (AWS), Google Cloud, and Microsoft Azure leading the way in offering flexible and secure services. Unlike traditional supercomputing clusters, where resources are allocated on an annual or bi-annual basis following rigorous scientific planning, cloud computing offers greater flexibility in resource allocation. Despite potentially higher associated costs, cloud computing provides researchers with an alternative that is well-suited to dynamic project requirements. With the support of AWS specialists, we successfully deployed a cloud-based supercomputer, utilizing Slurm for job submission management, as depicted in Fig. 8. The architecture of this supercomputer comprised a Slurm head node based on a C5 large instance, spot or on-demand Slurm compute nodes featuring Nvidia GPUs (P3 or P4D instances), customized Amazon Machine Images (AMIs), and essential libraries pre-installed. Additionally, a dedicated post-processing server equipped to execute Python-based scripts was included. Notably, all instances were interconnected through a shared Elastic File System (EFS). The distributed nature of the Adaptive Sampling strategy aligns seamlessly with cloud computing, as it operates without the need for synchronization during molecular dynamics trajectories. Such inherent adaptability renders the strategy agnostic to the specific computing resources it utilizes, making it well-suited for deployment within cloud computing environments.

**Fig. 8.**
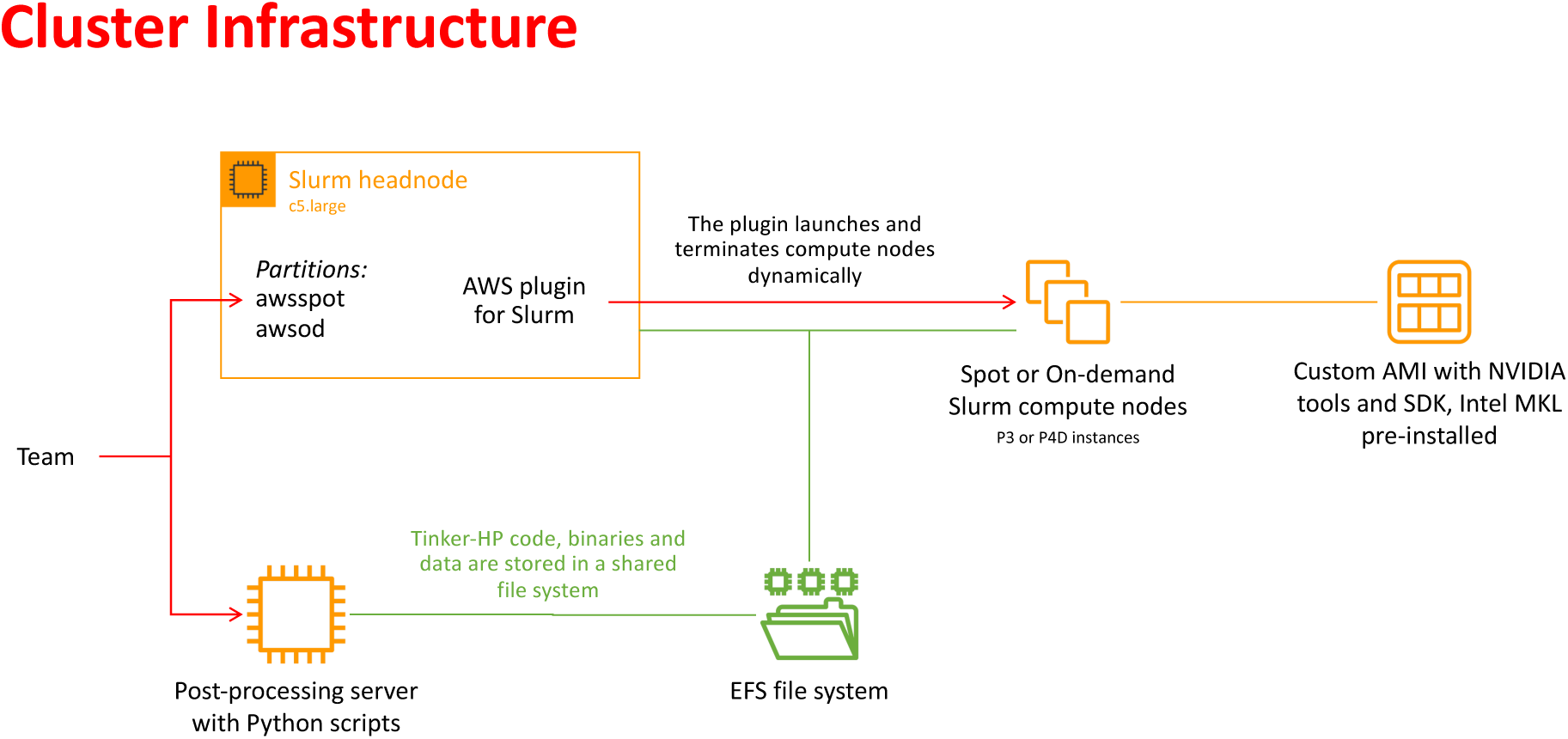
Schematic representation of the cloud-based virtual cluster used for this study, enabling the use of awsspot and awsod (on demand) partitions and leveraging a shared elastic file system.

### Statistical Analysis

#### Principal Component Analysis

The PCA analysis described in Results section was performed using Python libraries such as NumPy (*73*), MDTraj (*74*), Matplotlib (*33*), Scikit-learn (*75*), and SciPy (*76*).

Initially, we obtained trajectories of the spike protein from simulations available on the Amaro Lab website (*32*). Specifically, these trajectories were sourced from the work of Casalino et al. (*8*). The protein topology and scaffold residues were defined based on the Amaro PDB files corresponding to both open and closed states. Subsequently, the spike protein trajectories were aligned to the central scaffold residues, with a focus on the C*_α_* atom indices. Next, we applied PCA to the aligned trajectories to extract the dominant modes of motion exhibited by the spike protein. The PCA analysis was conducted using the Scikit-learn library, allowing us to compute the resulting principal components (PCs) and their corresponding explained variance ratios. The PCA coordinates were then saved for subsequent projection of our trajectories onto the obtained PCA space. To further characterize the distribution of conformations in the PCA space, kernel density estimation (KDE) was employed. Specifically, KDE was applied to each dimension of the PCA scores using the Gaussian KDE function from the SciPy library, facilitating the estimation of the probability density function of the data, providing valuable insights into the spatial distribution of spike protein conformations. The overall density at each point in the PCA space was calculated by multiplying the densities along each dimension, after which the density values were normalized. These normalized density values were utilized to generate a heatmap-style PCA plot, where each point represented a distinct spike protein conformation. The color of each point on the plot corresponded to the normalized density at that location, offering visual cues regarding the density distribution across the PCA space. The resulting plot was visualized using Matplotlib, and a color bar was included to indicate the density scale.

#### Root Mean Square Fluctuation

The Root Mean Square Fluctuation (RMSF) of protein C*_α_* atoms was computed to assess the dynamic behavior of the SARS-CoV-2 spike protein. Utilizing the Visual Molecular Dynamics (VMD) software package (*51*), the total number of frames in the molecular system was determined. The RMSF calculation was then performed using the **measure rmsf** utility available in VMD, which generated RMSF values for each C*_α_* atom over the trajectory frames.

#### Hydrogen Bond and Salt Bridge Analysis

The hydrogen bond and salt bridge analysis were conducted using VMD software environment (*51*) throughout the concatenated simulation trajectories for both open and closed states. For the hydrogen bond analysis, hydrogen bonds between protein residues were identified based on geometric criteria such as donor-acceptor distance (3 Å) and hydrogen-donor-acceptor angle (20°). Similarly, for the salt bridge analysis, interactions between positively and negatively charged protein residues were examined with an oxygen-nitrogen distance cut-off of 3.2 Å.

#### Solvent-Accessible Surface Area

The solvent-accessible surface area (SASA) of glycans was computed to investigate their solvent exposure throughout the simulation trajectory. The calculation was performed within the VMD software environment (*51*). Initially, a list of glycan segment names was defined to facilitate the selection process. The total number of frames in the trajectory was determined to iterate over each frame and over each glycan segment considering only its non-hydrogen atoms during the calculation. The SASA of the selected segment, excluding hydrogen atoms, was calculated using the **measure sasa** utility available in VMD with a probe radius of 1.4 Å. The SASA values obtained for each frame were accumulated to compute the total SASA for the segment. Subsequently, the average SASA for the segment across all frames was determined by dividing the total SASA by the number of frames.

#### Center-of-Mass Distances between Protein and Glycan Residues

The selection of protein domains and corresponding glycans was based on Table S1 in Casalino et al. (*8*). To compute the distances between these selected residues, the trajectory was iterated over. Utilizing VMD **measure center** utility, the center-of-mass (COM) coordinates for each selected protein and glycan group were determined, followed by the computation of the distance between their respective COMs.

#### Dynamic Cross-Correlation Function

The analysis of dynamic cross-correlations between protein residues and glycan segments was conducted using a combination of Python libraries, including NumPy (*73*), Mat-plotlib (*33*), SciPy (*76*), MDAnalysis (*77, 78*), and Seaborn (*79*). The protein topology and trajectory were processed using the MDAnalysis library within Python. The protein residues and glycan segments of interest were selected based on predefined residue numbers and segment names. The center of mass (COM) for both the selected protein residues and glycan segments was calculated for each frame of the trajectory. The COM values were stored in NumPy arrays for further analysis. Using the **correlate2d** function from the SciPy library, the dynamic cross-correlation matrix was computed between each pair of residues and glycan segments throughout the trajectory. To ensure consistent visualization, the dynamic cross-correlation matrix was normalized using a min-max normalization function. The intensity of each cell in the heatmap indicates the strength of the correlation between the corresponding residue and glycan segment.

#### 0.0.1 Radial Distance Analysis of Glycans

To investigate the radial distribution of the glycan shield, we calculated the radial distances of glycan COM relative to the z-axis across all frames of the simulation (see Fig. 4B). For each frame, we selected all glycan residues across specified segments (G1 to G70), excluding hydrogen atoms. The COM of the selected glycan residues was then computed within VMD **measure center** utility. The radial distance of the glycan COM from the z-axis was determined using the formula 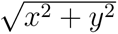, where x and y are the coordinates of the COM in the xy-plane.

#### Radial Distribution Analysis. Coordination Number. Pair Distribution Analysis

We leveraged Radial Distribution Function (RDF) analysis to unravel the intricate distribution patterns of interacting oxygen atoms across diverse molecular interactions, notably within protein-water and glycan-water interfaces. Similar to SASA, the RDF analysis was performed within the VMD software environment (*51,80*). A bin size of 0.1 Å and a total radial distance of 10 Å were used. The selection of oxygen atoms from water, glycan, and protein was customized based on the specific aims of our investigation discussed in the main text. Besides, to evaluate the number of water molecules in the proximity to the selected protein and glycan residues, we meticulously evaluated coordination numbers through the integration of RDFs, providing nuanced insights into the degree of atomic coordination within the molecular system under scrutiny. Additionally, we conducted localized analyses akin to RDF, employing pair distribution function (PDF) calculations to delve deeper into the distribution profiles of specific atom pairs within the same tool.

#### Polarizable Water Indication

In the initial stage, we compiled lists of water molecules exhibiting high dipole moments (> 2.8 D) for each frame in every MD step of open-state simulations. Through careful observation, we then visually (i.e., by means of VMD software (*51*)) identified regions where these molecules clustered most frequently, particularly focusing on their proximity to specific protein residues. Subsequently, we established a threshold of 10 Å to investigate changes in solvent dynamics near the associated protein residues along all the accumulated trajectories. For each water molecule proximity to the target protein and glycan residues within each micro-iteration, we calculated the average dipole moment, average distance from the protein residue, and occupancy percentage throughout the simulation. The average dipole moment was computed by summing the dipole moments of all water molecules within the defined region and dividing them by the total count of water molecules. Similarly, the average distance from the protein and glycan residues was determined by calculating the distance between each water molecule and the COM of the target residues. The occupancy percentage was calculated to quantify the extent to which water molecules occupied the defined region around the selected residues (at a distance less than 5 Å). In addition, each occupancy percentage was weighted based on the weights obtained from the density PCA of our conformational sampling method, ensuring that the contribution of each occupancy value to the overall analysis was proportional to its significance in the conformational space. The weighted occupancy for each water molecule was calculated by dividing the occupancy percentage by the total weight obtained from the density PCA. The obtained data was further filtered within the criteria defined in the main text in the Results section, ensuring that only interactions meeting specific thresholds were considered for subsequent analysis.

Given the absence of water molecules with high dipole moments (i.e., exceeding 2.8 D) in the closed-state simulations, we opted to analyze the average dipole moments of protein and glycan atoms known to undergo the previously described phases in the open configuration. Additionally, we calculated the average dipole moments of oxygen atoms of water molecules proximal to these residues to glean insights into their polarizability dynamics and compared them to those of the open state, thus offering a comprehensive understanding of solvent-protein interactions across various simulation conditions.

#### Dipole Moment Distribution Analysis

The dipole moment distribution analysis involved extracting dipole moment data from simulation output files and visualizing their distribution. Firstly, all relevant data files containing dipole moment information were located within the specified directory corresponding to each of the defined phases of water interaction. Next, the dipole moments were extracted from each file. The average dipole moment and its standard deviation were calculated across all frames of the simulation trajectories. Subsequently, a histogram with a kernel density estimate was plotted using the Seaborn library (*79*). This visualization provided an overview of the dipole moment distribution, with the x-axis representing the dipole moment values and the y-axis representing probability density. Additionally, vertical lines indicating the mean, median, and mode of the dipole moments were overlaid on the histogram, offering further insights into the distribution characteristics.

## Author contributions

Conceptualization: MB, LL, JPP Methodology: OA, CL, LL, PR, JPP Investigation: MB, LL Visualization: MB Supervision: LL, JPP Writing—original draft: MB Writing—review & editing: LL, JPP, CL, PR

## Data and materials availability

All the data used in the analyses are available in the paper and/or the Supplementary Materials. Further information and requests should be directed to and will be fulfilled by the lead contact, JPP.

## Supplementary Materials

This PDF file includes:

Figs. S1 to S13

Legend for data S1 to S4

References 1 to 4

**Other Supplementary Material for this manuscript includes the following:**

Data S1 to S4 Movie S1

**Fig. S1.**
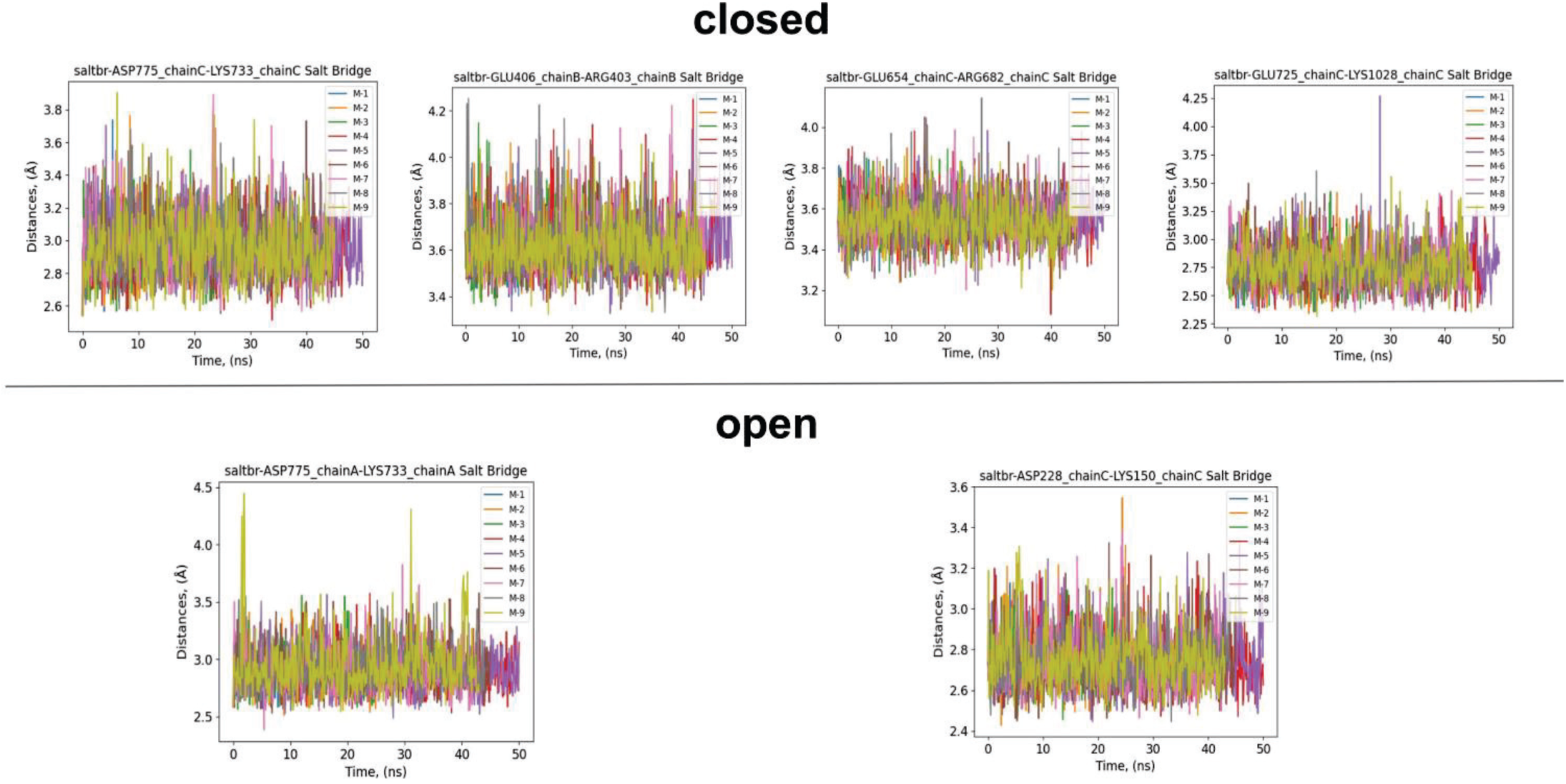
Stable salt bridges in the SARS-CoV-2 spike protein structure, shown for the closed (upper panel) and open (lower panel) conformations across all macro-iterative simulations. These simulations were conducted using the polarizable AMOEBA force field and density-driven conformational sampling, as detailed in the Methods section of the main text (M-refers to the macroiteration). Chains A, B, and C represent the trimeric configuration of the protein.

**Fig. S2.**
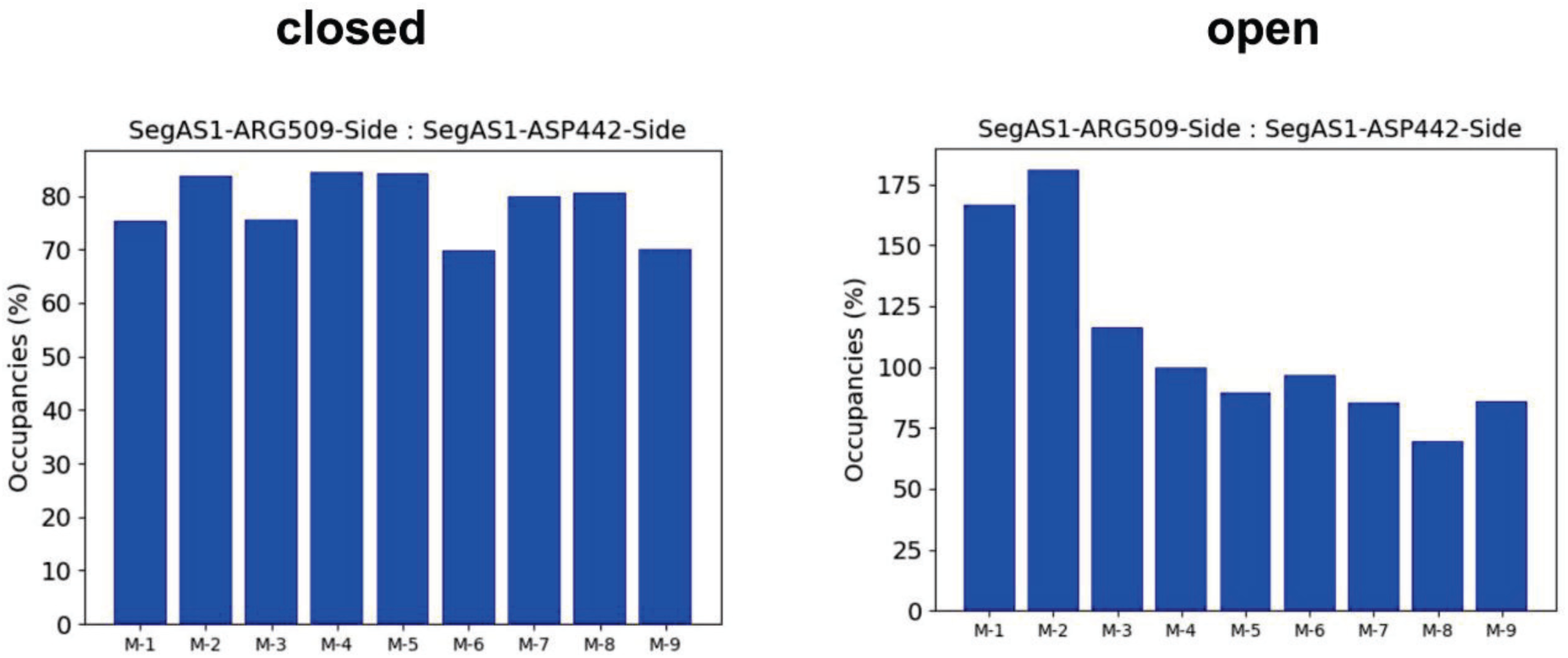
Common stable inner hydrogen bonds found in the RBD domain of both closed and open states along all the macroiterative simulations.

**Fig. S3.**
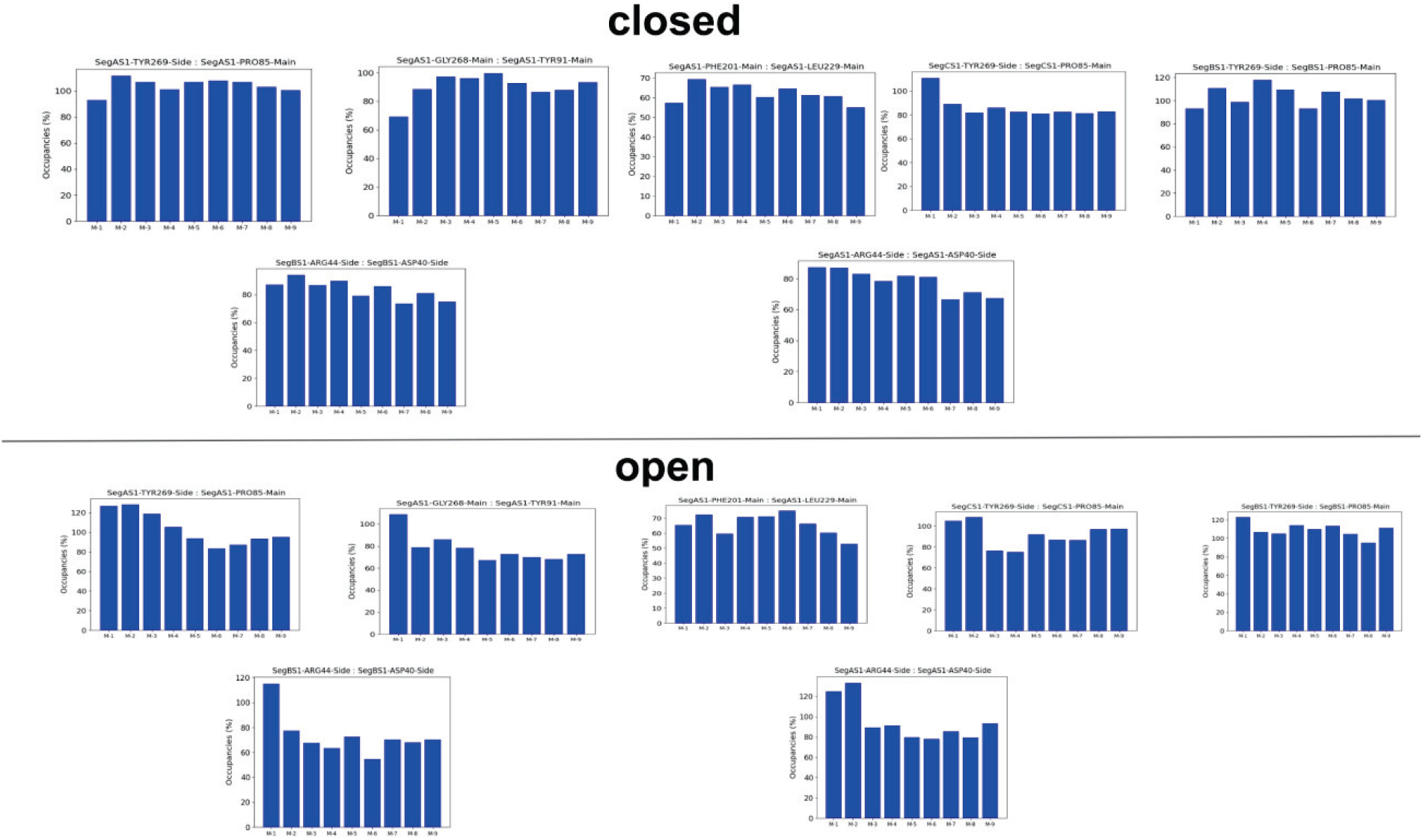
Common stable inner hydrogen bonds found in the NTD domain of both closed and open states along all the macroiterative simulations.

**Fig. S4.**
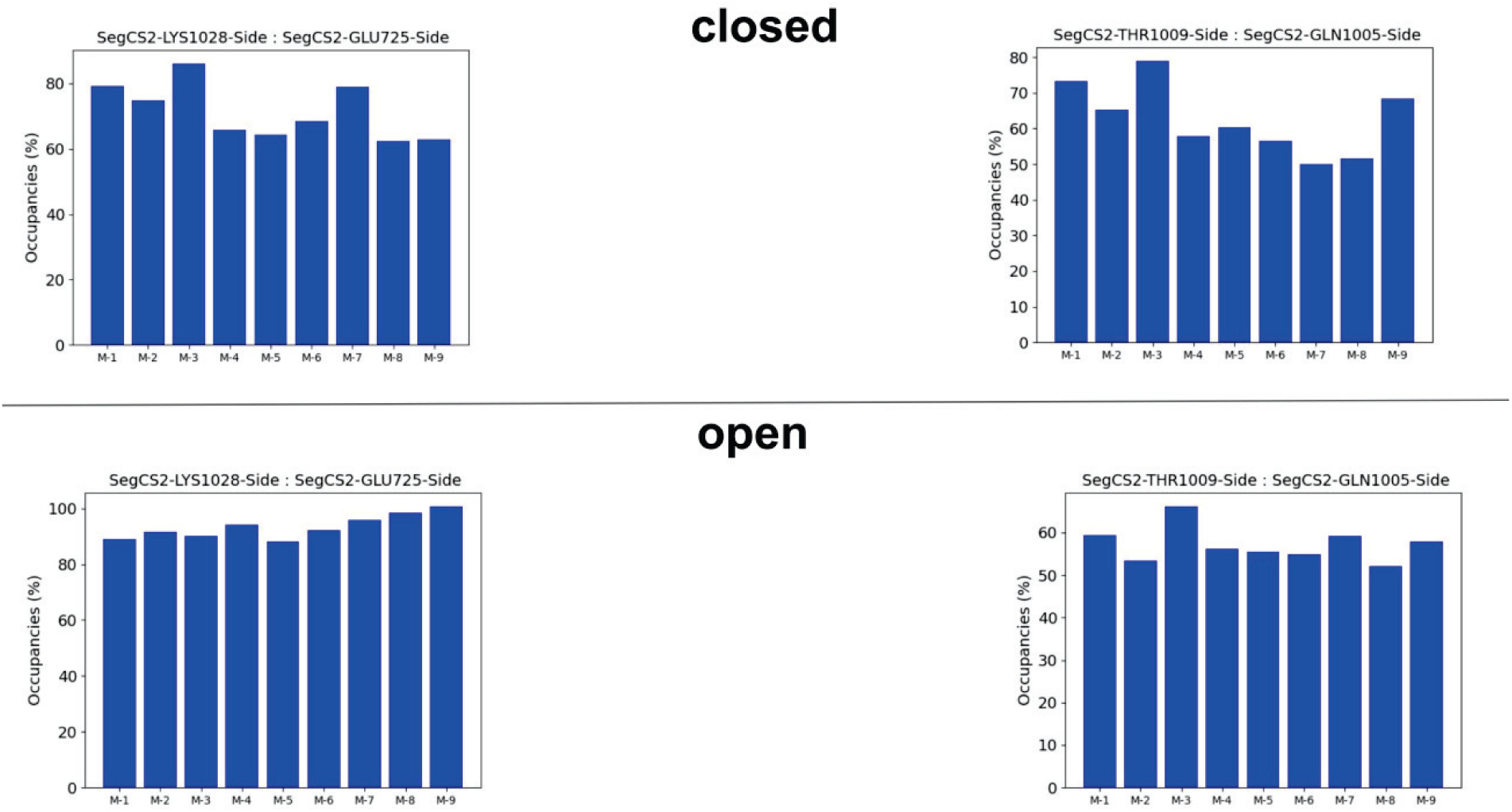
Common stable inner hydrogen bonds found in the CH domain of both closed and open states along all the macroiterative simulations.

**Fig. S5.**
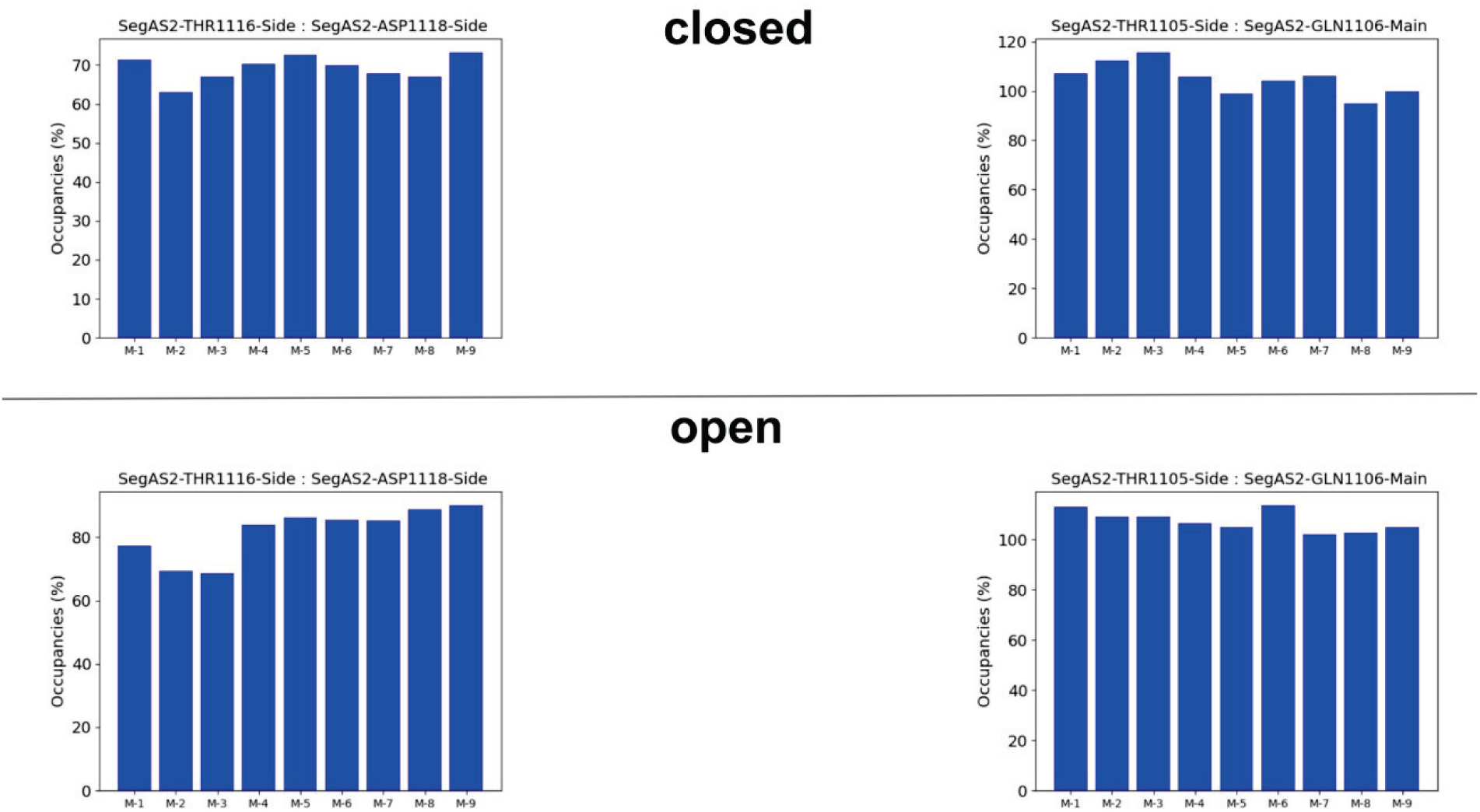
Common stable inner hydrogen bonds found in the CD domain of both closed and open states along all the macroiterative simulations.

**Fig. S6.**
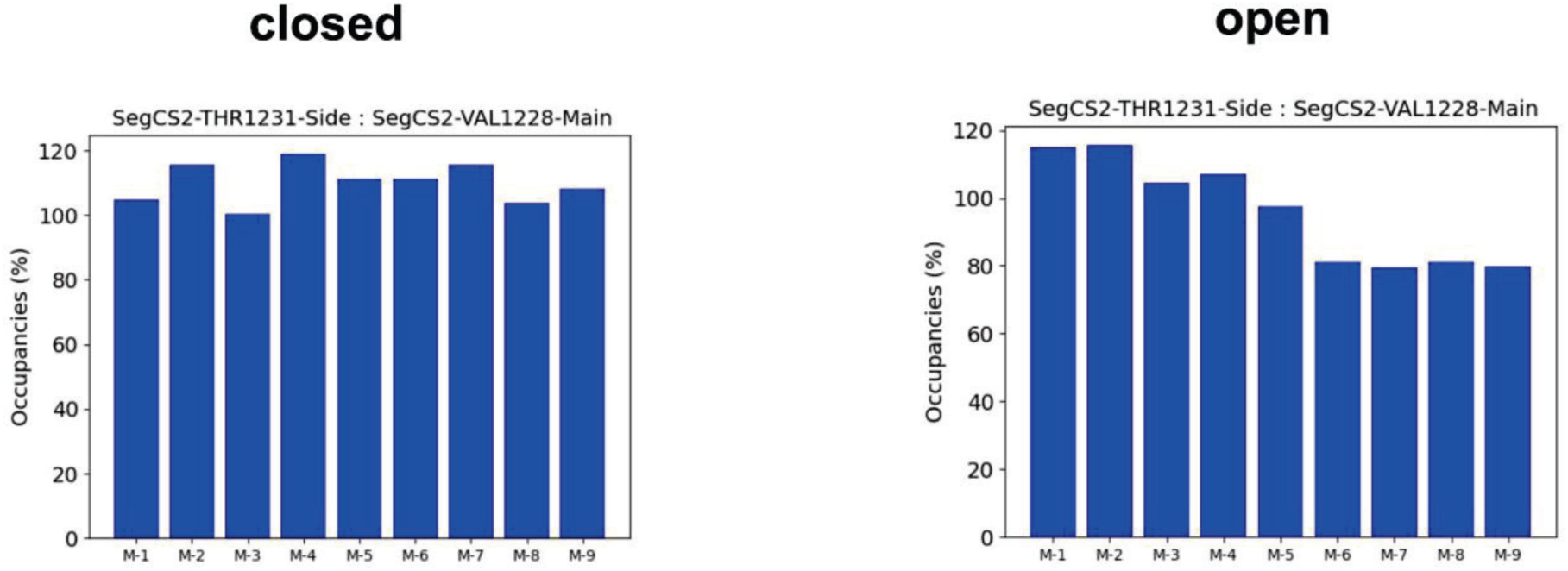
Common stable inner hydrogen bonds found in the TM domain of both closed and open states along all the macroiterative simulations.

**Fig. S7.**
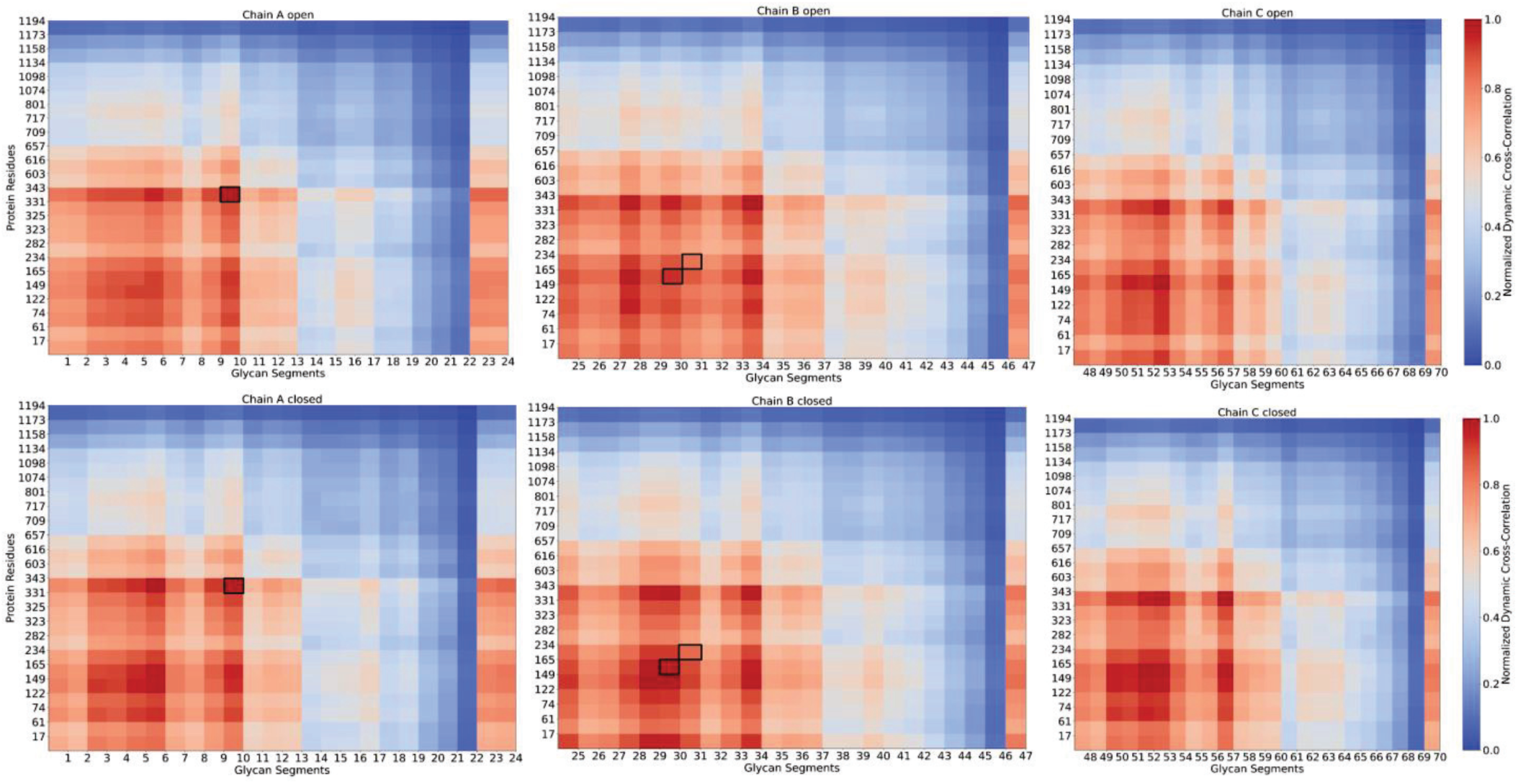
Normalized dynamic cross-correlation function depicting the dynamic interactions between protein residues of chains A, B, C and surrounding glycan segments across open (top) and closed (bottom) states of the SARS-CoV-2 viral structure. The color intensity represents the strength of correlation, ranging from 0 (low correlation, shown in blue) to 1 (high correlation, shown in red). The residues responsible for the glycan gating are contoured with rectangle black boxes.

**Fig. S8.**
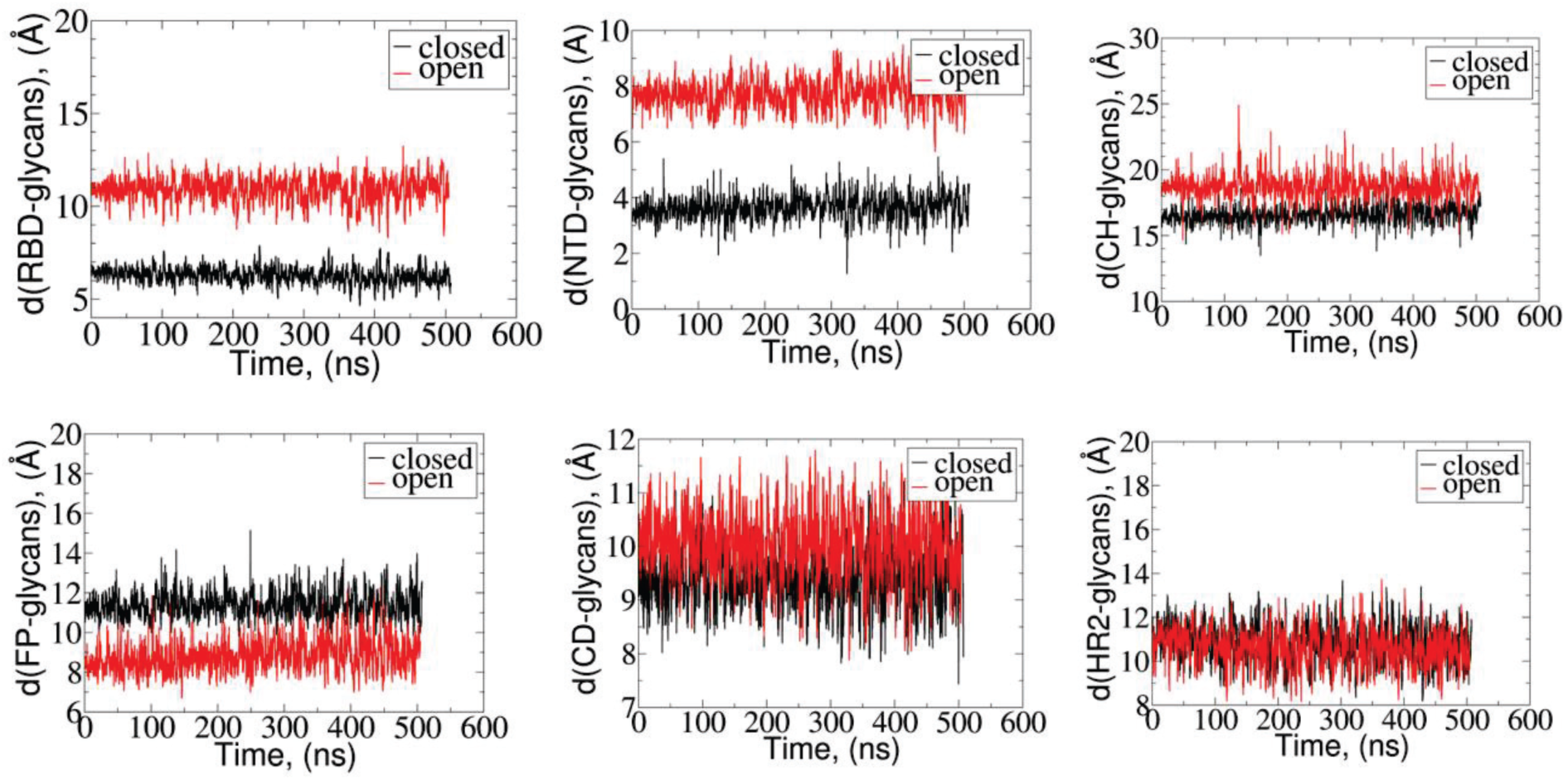
Center-of-mass (COM)-to-COM distances between protein domains and surrounded glycans along combined simulations from all iterations for open (red) and closed (red) states.

**Fig. S9.**
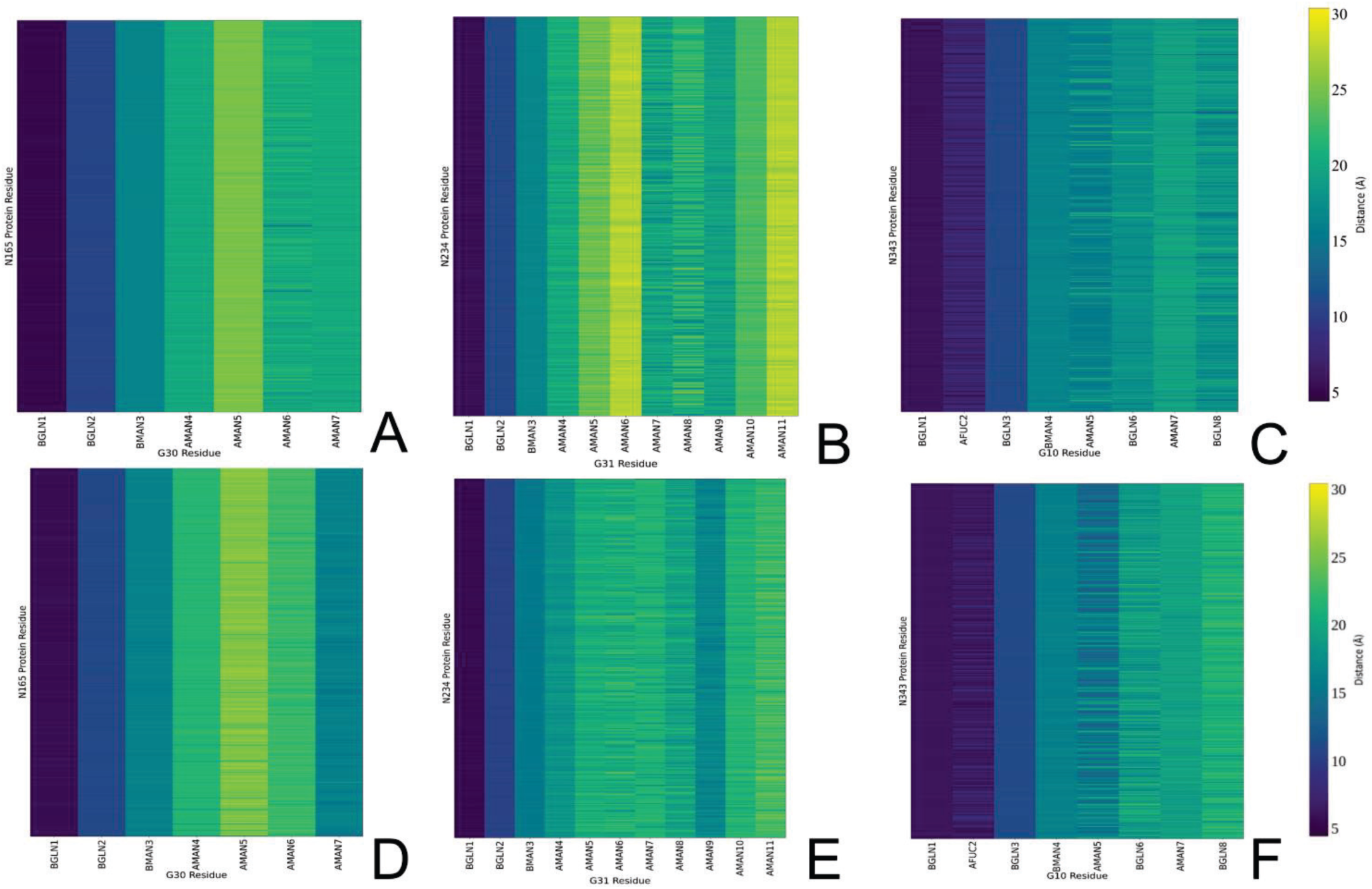
Contact maps of interactions between center-of-mass (COM) of N165, N234, and N343 of the protein and corresponding glycans G30, G31, and G10 in open (A, B, C) and closed (D, E, F) states. The corresponding COM of glycan residues are BMAN (β-D-mannose), AMAN (*α*-D-mannose), BGAL (β-D-galactose), BGLN (β-D-glucoseamine), ANE5 (α-D-neuraminic acid (also known as sialic acid)), AGAN (α-D-galactose), AFUC (α-L-fucose).

**Fig. S10.**
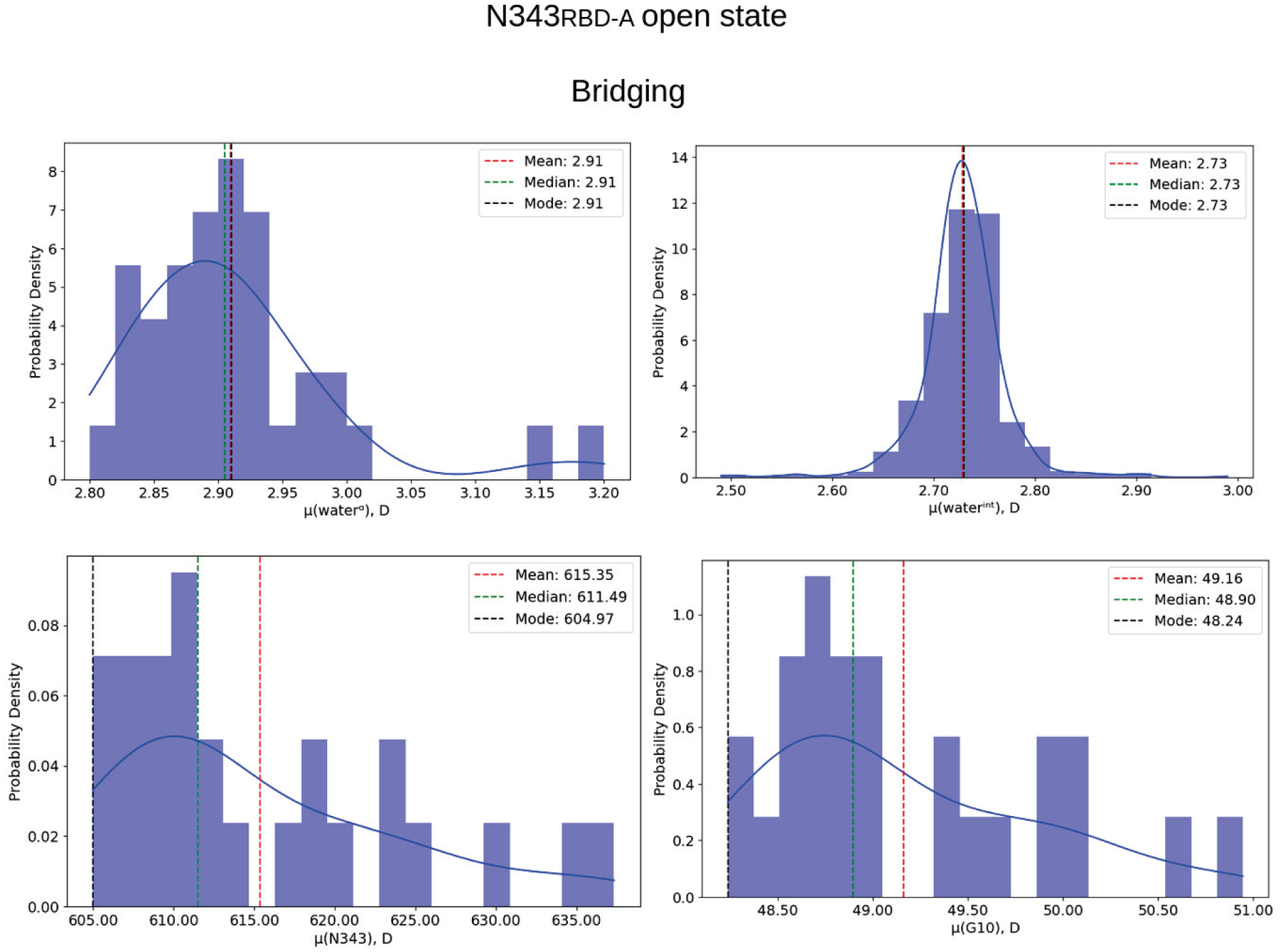
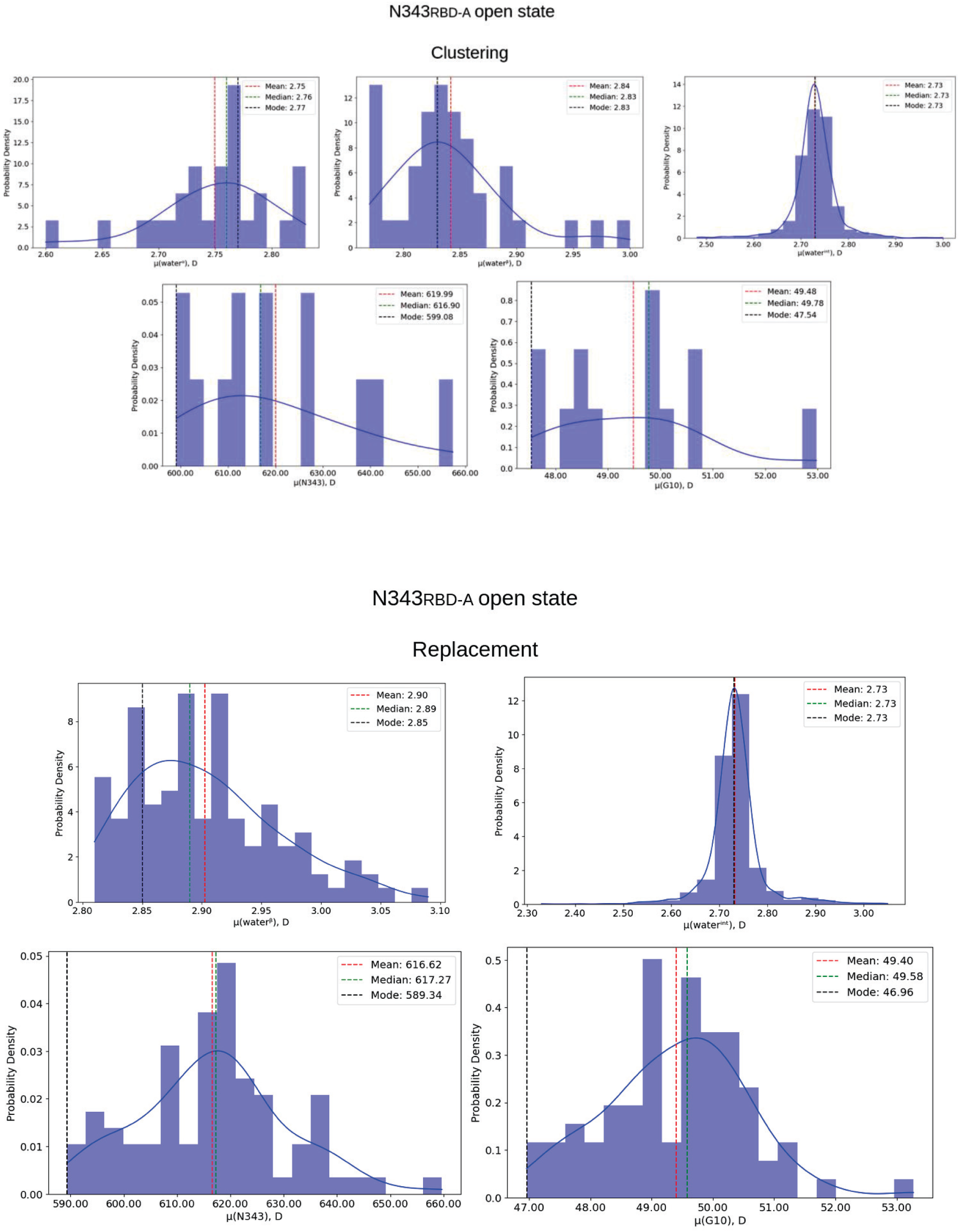
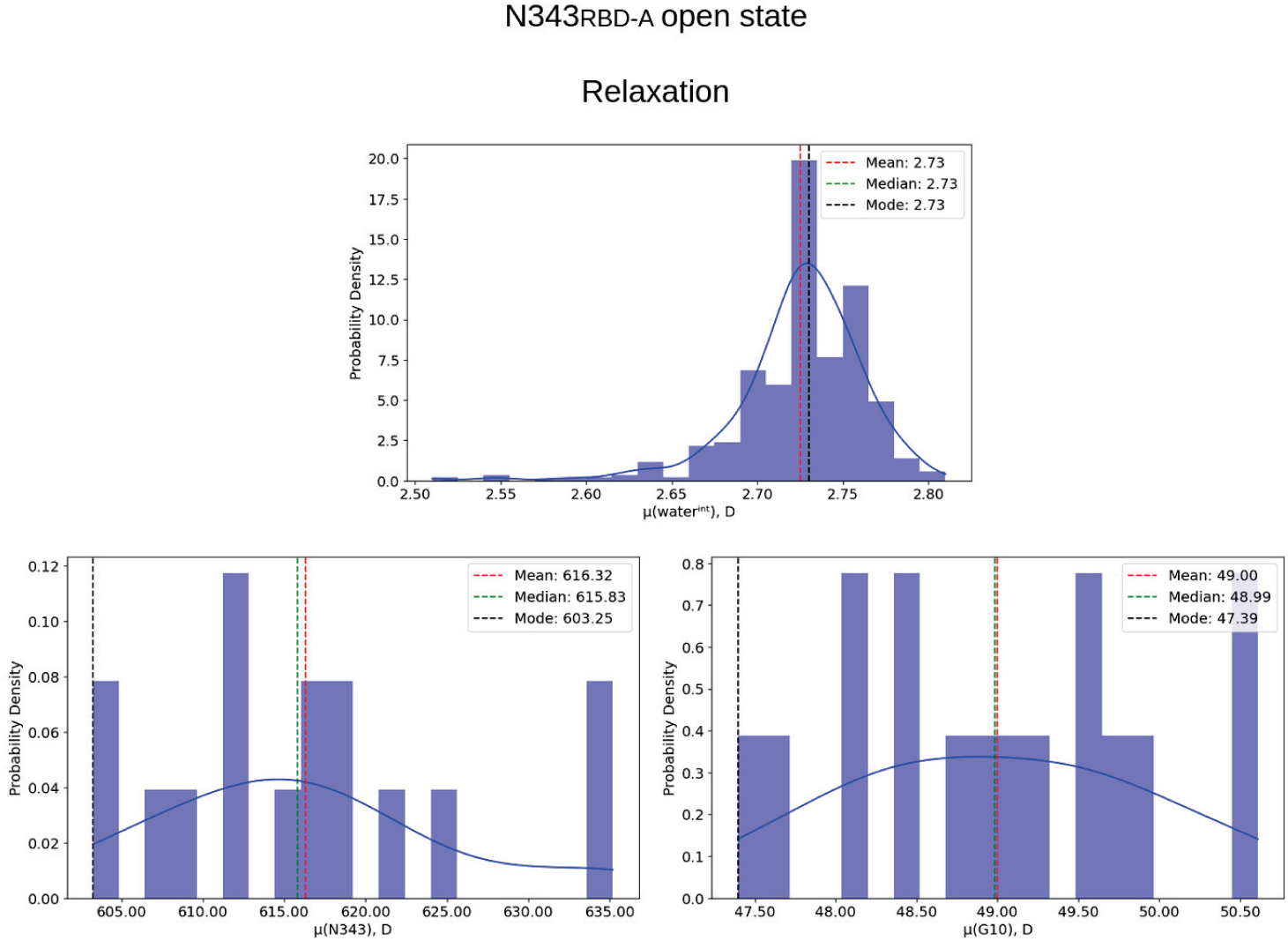
Distribution of dipole moments along each phase for the N343_RBD-A_ interaction pattern in the open state, including mode, mean, median values.

**Fig. S11.**
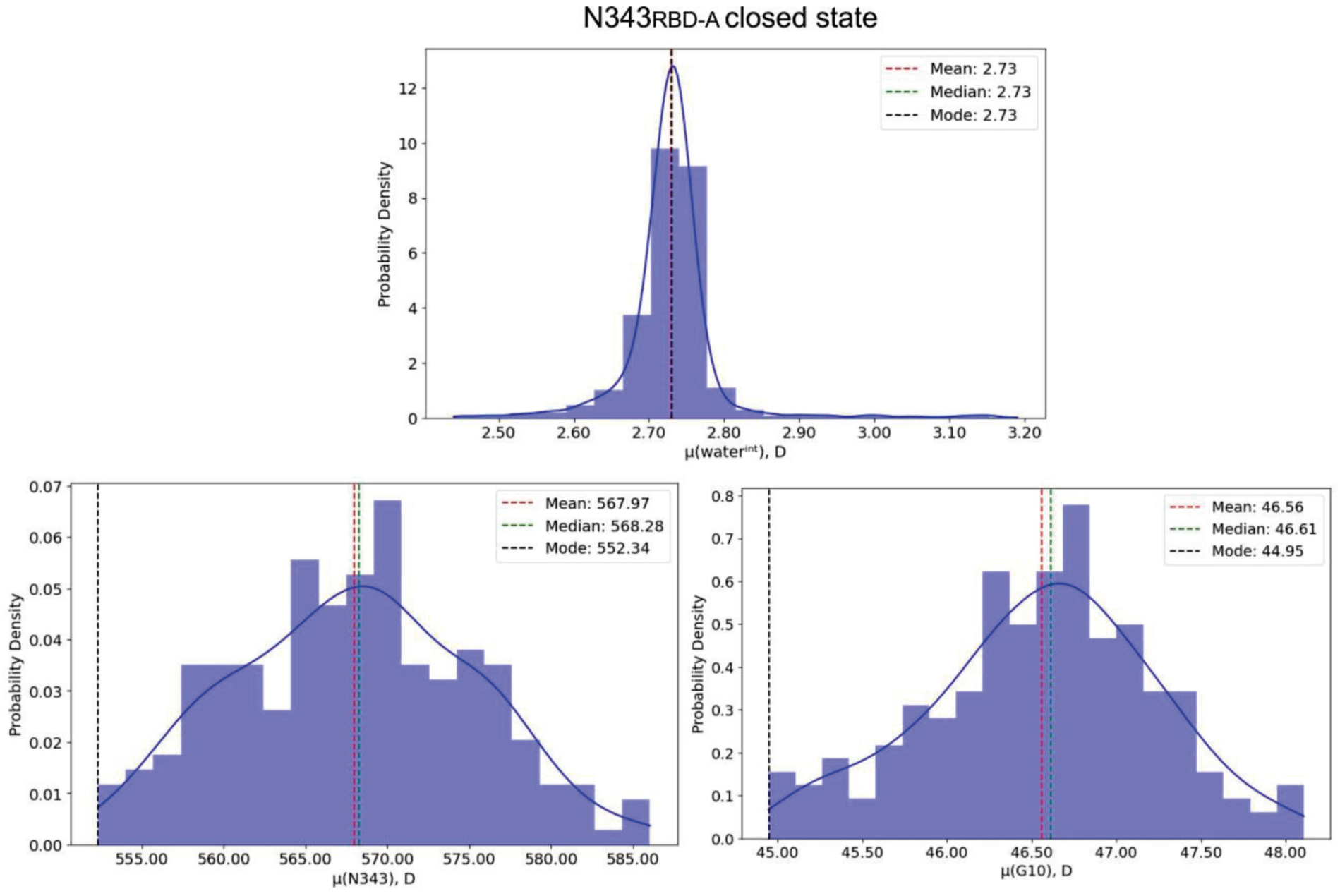
Distribution of dipole moments along each phase for the N343_RBD-A_ interaction pattern in the closed state, including mode, mean, median values.

**Fig. S12.**
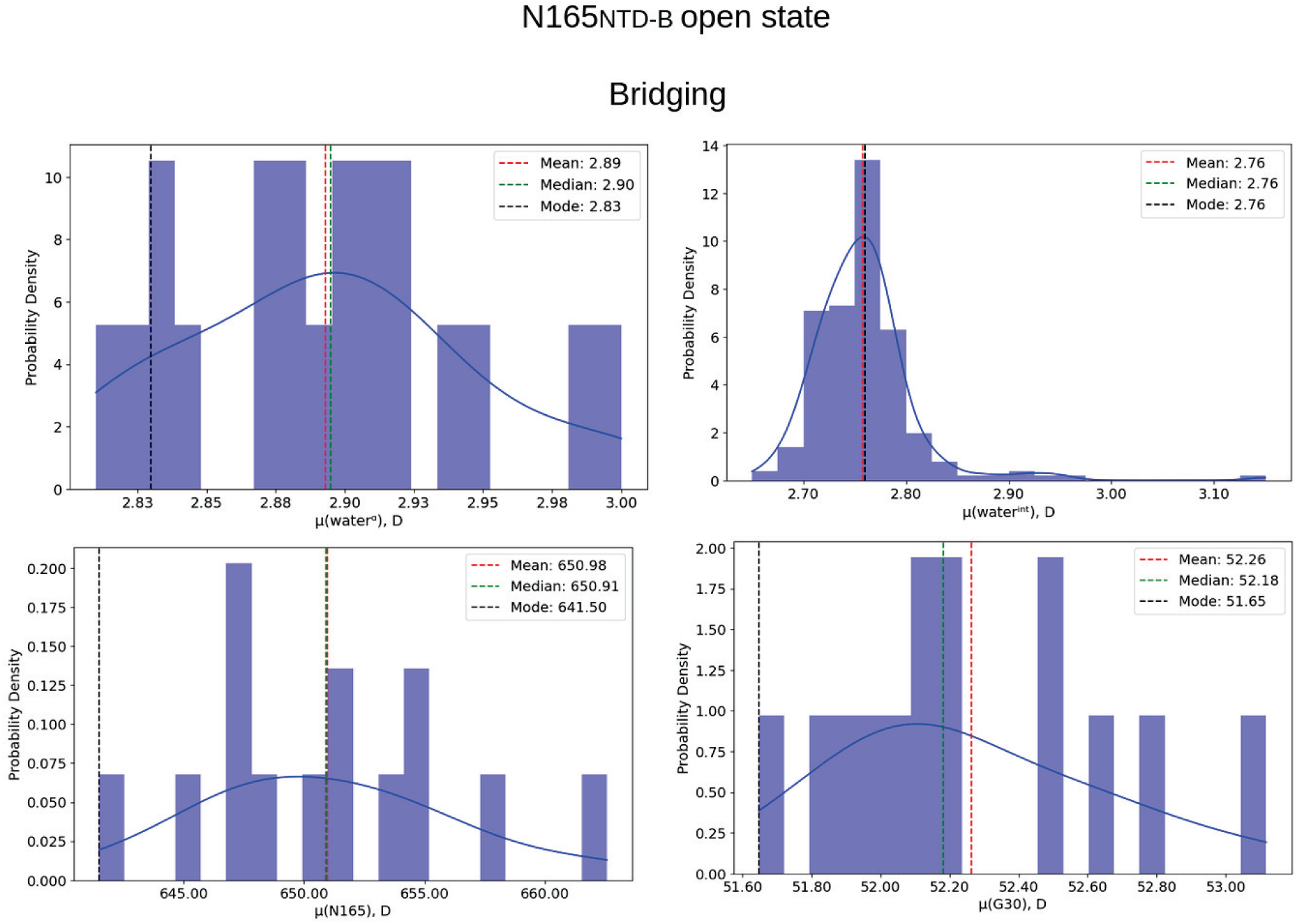
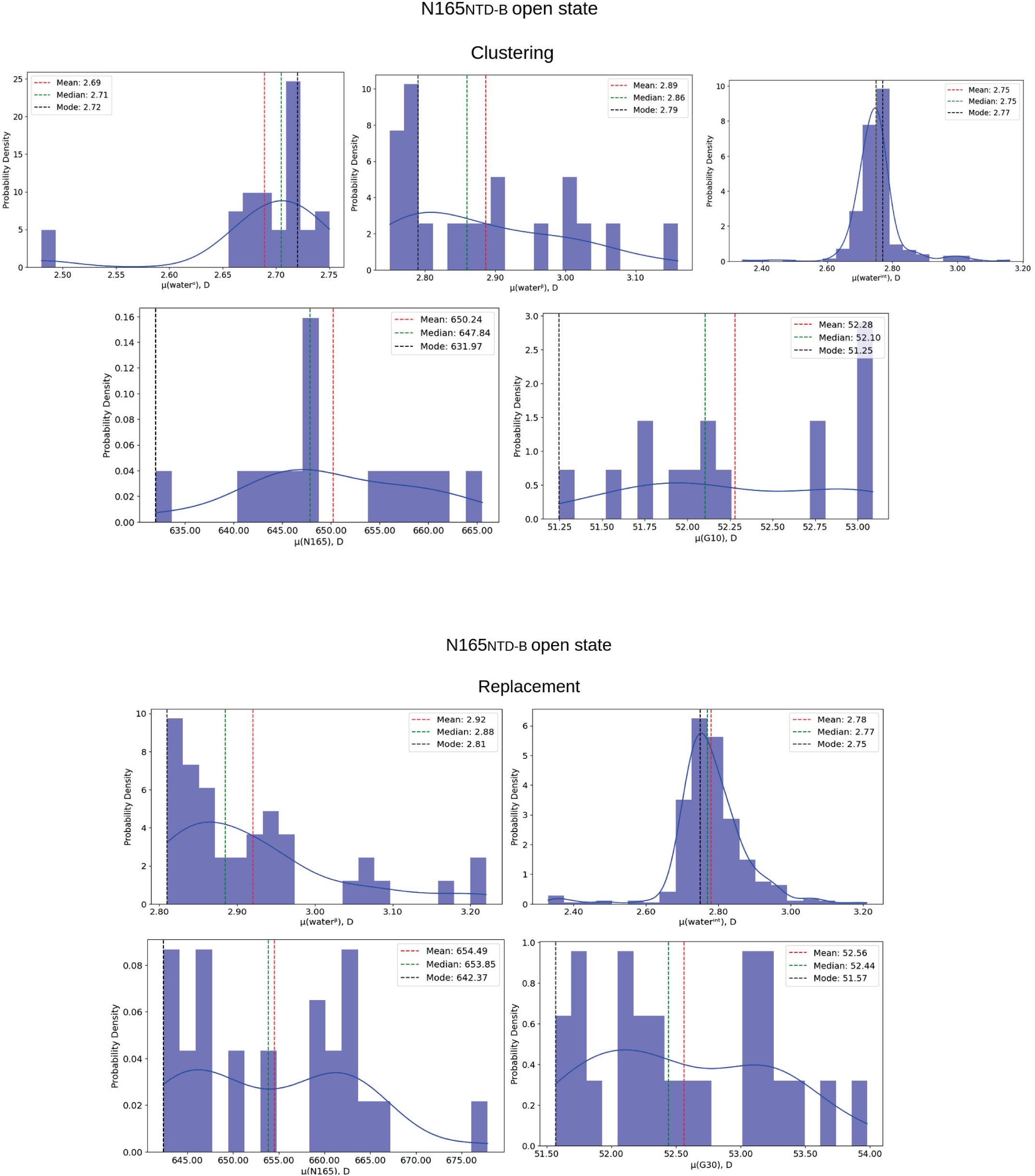
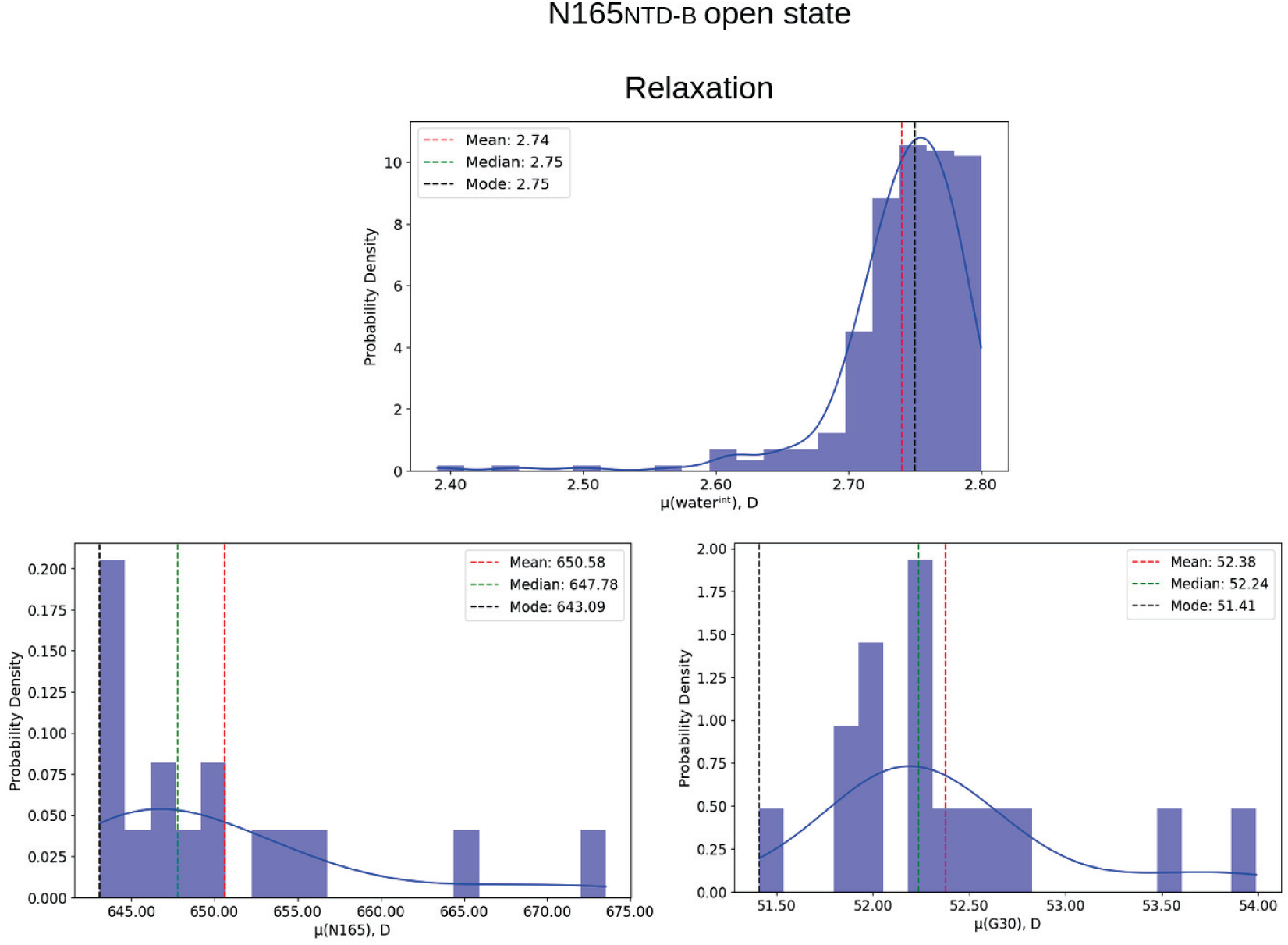
Distribution of dipole moments along each phase for the N165_NTD-B_ interaction pattern in the open state, including mode, mean, median values.

**Fig. S13.**
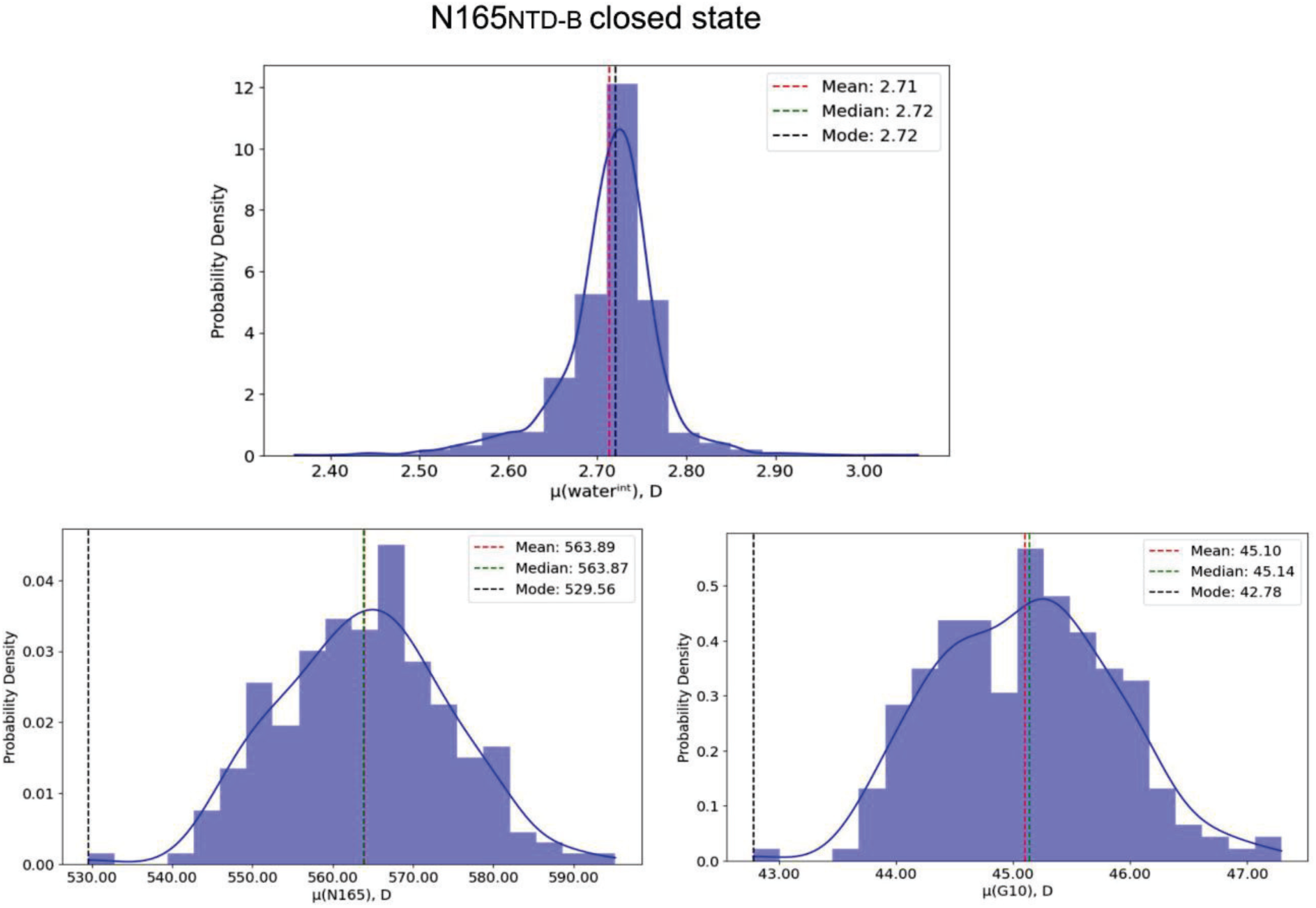
Distribution of dipole moments along each phase for the N165_NTD-B_ interaction pattern in the closed state, including mode, mean, median values.

## Parametrization details for glycans and lipid molecules

AMOEBA parameters for the glycan molecules, including BMAN (beta-D-mannose); AMAN (alpha-D-mannose), BGAL (beta-D-galactose), BGLN (beta-N-acetyl-D-glucosamine), ANE5 (N-acetyl-alpha-neuraminic acid), AGAN (N-acetyl-alpha-D-galactosamine), AFUC (alpha-L-fucose), and lipid molecules, including POPC (1-palmitoyl-2-oleoyl-sn-glycero-3-phosphocholine), POPI (1-palmitoyl-2-oleoyl-sn-glycero-3-phosphoinositol), POPS (1-palmitoyl-2-oleoyl-sn-glycero-3-phosphatidylserine), and cholesterol were derived using the Poltype2 automation tool discussed in the main text. For POPE (1-palmitoyl-2-oleoyl-phosphatidyl ethanolamine), the parameters were taken from reference (*1*) as well as the parameters in common for all lipids oleoyl motifs. The sn-glycero-3-phosphocholine (in POPC), sn-glycero-3-phosphoinositol (in POPI), and sn-glycero-3-phosphatidylserine (in POPS) parts and cholesterol were parametrized separately. Each input glycan residue and individual non-oleoyl lipid fragments (in **sdf** format) was first optimized at MP2/6-31G* level of theory. Optimized geometry then was used for two single-point calculations at MP2/6-311G** (low-level) and MP2/aug-cc-pvtz (high-level) of theory, respectively. The electron density of the low-level calculations was used to derive the atomic multipoles, by employing the distributed multipole analysis method in GDMA program (available in open-source library https://gitlab.com/anthonyjs/gdma) (*2*). These multipoles were further optimized by fitting to the electrostatic potential (ESP) generated by the high-level electron density. The valence and torsion parameters were matched to the existing database in Poltype2 by using SMARTS patterns (see https://www.daylight.com/dayhtml/doc/theory/theory.smarts.html) (*3*). For the remaining torsion parameters that were not found in SMARTS database, the dihedral angle around the rotatable bond was spined over 360 degrees at interval of 30 degree (12 datapoints for each dihedral angle). For the torsional angle in the glycan rings, 5 datapoints (perturbed from the original angle value by - 20, -10, 0, +10, +20 degrees). The geometry of each structure at a certain dihedral value was then optimized at PBE1PBE/6-31G level of theory by restraining the torsion angles of interest and followed by single point energy calculations at MP2/6-31+G* level of theory. The relative energy from the QM calculations then were targeted to fit the torsional parameters. All the QM calculations were performed using Gaussian 09 package (*4*). The **poledit.x** executable within Tinker software was used to convert the multipole values from GDMA output (in global frame) to Tinker format (in its local frame definition). The **potential.x** executable in Tinker was used to refine the multipoles by fitting to high-level ESP generated by QM. Parameter files and all the torsion fit plots have been included in the Supplementary Material with several exceptions: (1) ANE5 since all the torsional parameters have been matched from database and for AGAN the acetyl-amine fragment along with galactose core were merged from BGAL and BGLN parametrization outputs and (2) POPE parameters. Additionally, we included the final parameter file and the corresponding Tinker **xyz** structure files for both closed and open states.

Legend for **Data S1**: Calculated inner protein hydrogen bonds for open and closed states.

Legend for **Data S2**: Parametrization details for glycans, lipids.

Legend for **Data S3**: Structure files for closed state.

Legend for **Data S4**: Structure files for open state.

Legend for **Movie S1**: Artistical visualization of the open and closed state systems used in our study.

